# Elevated energy costs of biomass production in mitochondrial-respiration deficient *Saccharomyces cerevisiae*

**DOI:** 10.1101/2022.11.15.516572

**Authors:** Pranas Grigaitis, Samira L. van den Bogaard, Bas Teusink

## Abstract

Microbial growth requires energy for maintaining the existing cells and producing components for the new ones. Microbes therefore invest a considerable amount of their resources into proteins needed for energy harvesting. Growth in different environments is associated with different energy demands for growth of yeast *Saccharomyces cerevisiae*, although the cross-condition differences remain poorly characterized. Furthermore, a direct comparison of the energy costs for the biosynthesis of the new biomass across conditions is not feasible experimentally; computational models, on the contrary, allow comparing the optimal metabolic strategies and quantify the respective costs of energy and nutrients. Thus in this study, we used a resource allocation model of *S. cerevisiae* to compare the optimal metabolic strategies between different conditions. We found that *S. cerevisiae* with respiratory-impaired mitochondria required additional energetic investments for growth, while growth on amino acid-rich media was not affected. Amino acid supplementation in anaerobic conditions also was predicted to rescue the growth reduction in mitochondrial respiratory shuttle-deficient mutants of *S. cerevisiae*. Collectively, these results point to elevated costs of resolving the redox imbalance caused by *de novo* biosynthesis of amino acids in mitochondria. To sum up, our study provides an example of how resource allocation modeling can be used to address and suggest explanations to open questions in microbial physiology.

## Introduction

Energy turnover is central to life as we know it: living cells use energy for many of their functions, such as to maintain chemical gradients across membranes, to polymerize macromolecules (Lahtvee et al., 2017; Verduyn et al., 1990a), to perform mechanical work, or to condition unfavorable environments (e.g. to neutralize toxic compounds). As a result, cells invest a substantial amount of available resources into energy-harvesting pathways: quantitative proteomics measurements of budding yeast *Saccharomyces cerevisiae* grown in aerobic, glucose-limited chemostats suggested that up to 1/3 of the total cell proteome is allocated to energy generation pathways (Elsemman et al., 2022).

The steady-state energy turnover is described as an equal rate of sum energy extraction from nutrients and investment into growth. *S. cerevisiae* intensively ferments glucose into ethanol in glucose-excess conditions (Blank and Sauer, 2004), yet, fluctuations of nutrient levels are a frequent phenomenon in both natural and biotechnologically-relevant environments (Haringa et al., 2016). There, oscillations of glucose availability, for instance, are handled by anticipatory protein expression (Grigaitis and Teusink, 2022; van den Brink et al., 2008). Thus the cells in glucose-scarce conditions are indeed primed to consume the incoming flux of nutrients and store the released energy in the high-energy bonds of the ATP molecule.

The other side of the balance, ATP investment into the biosynthesis of new cell components, tends to be more dynamic. First, the cell composition of *S. cerevisiae* can greatly vary across conditions, even for the same growth-limiting substrate as its availability changes (Canelas et al., 2011; Lange and Heijnen, 2001). Second, external factors (e.g. presence of oxygen) can dictate the availability of some of the biosynthesis routes.

Costs of producing new cell biomass from nutrients can be computed from the biosynthesis routes of major cell components: proteins, nucleic acids, lipids, and carbohydrates. Computational models, namely, genome-scale metabolic models, can be of great help in this endeavor. A genome-scale metabolic model (GEM) is a compendium of the reactions that can happen in an organism based on its genome sequence (Lu et al., 2019). To the day, GEMs are the most extensive tools to approximate biochemistry in a fine-grained manner at large-scale and have a diverse range of applications (Somerville et al., 2022). Extended genome-scale models which also address the protein costs of running metabolic reactions can aid to identify the most optimal metabolic strategies in terms of allocation of cellular resources (Elsemman et al., 2022).

Prediction of condition-dependent metabolic strategies and quantification of energy- and nutrient costs for biomass production can explain metabolic phenotypes and offer targets to optimize industrial processes. In this study, we present an updated resource allocation model of *S. cerevisiae* which we used to identify and metabolic strategies of yeast across conditions and genotypes. We found that growth with impaired- or inactive mitochondria in chemically defined minimal media requires additional energetic and proteomic investments. The model predictions pointed to additional costs of resolving the surplus of mitochondrial redox equivalents accumulated due to biosynthetic processes in mitochondria. Overall, we argue that the computational analysis of microbial metabolic strategies is a powerful approach to deepen understanding of microbial physiology.

## Results

### The updated proteome-constrained model of *S. cerevisiae, pcYeast8*

We improved our previously published version of the protein constrained model of yeast (*pcYeast7*.*6*; (Elsemman et al., 2022)) and created an updated version, *pcYeast8*. In this version, we also curated the *Yeast8* metabolic model (Lu et al., 2019): the major improvements were the reconstruction of new pathways related to lipoic acid synthesis, curation of the glycerol:*H*^+^ symport reaction, and others (Supplementary Notes).

We first used the model to revisit our earlier analysis of growth in aerobic glucose-limited chemostats (Figure 1a-c). A major improvement to the *pcYeast7*.*6* model is the correct prediction of glycerol excretion at high growth rates (Figure 1a). Furthermore, the respiratory quotient 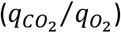, which represents the ratio of flux through respiration versus fermentation, and the biomass yield of glucose (*Y*_*X/S*_) matches the experimental data well (Figure 1b-c). Active proteome constraints, which give a detailed overview of the growth-limiting constraints at every simulation step, remained as previously reported. The model also predicts proteome constraints which actively limit growth at every simulation step: at low specific growth rates, and before the critical dilution rate, glucose transport capacity actively limited growth, to be followed by the mitochondrial capacity constraint, and after *µ =* 0*.3*5 *h*^−1^, limitation in the cytosolic proteome capacity (Figure 1d). At maximal growth rate (glucose excess conditions), the only active proteome constraint was the cytosolic proteome capacity. The sequence of constraints hit as a function of growth rate was consistent with the earlier model version we published (Elsemman et al., 2022).

**Figure 1.**
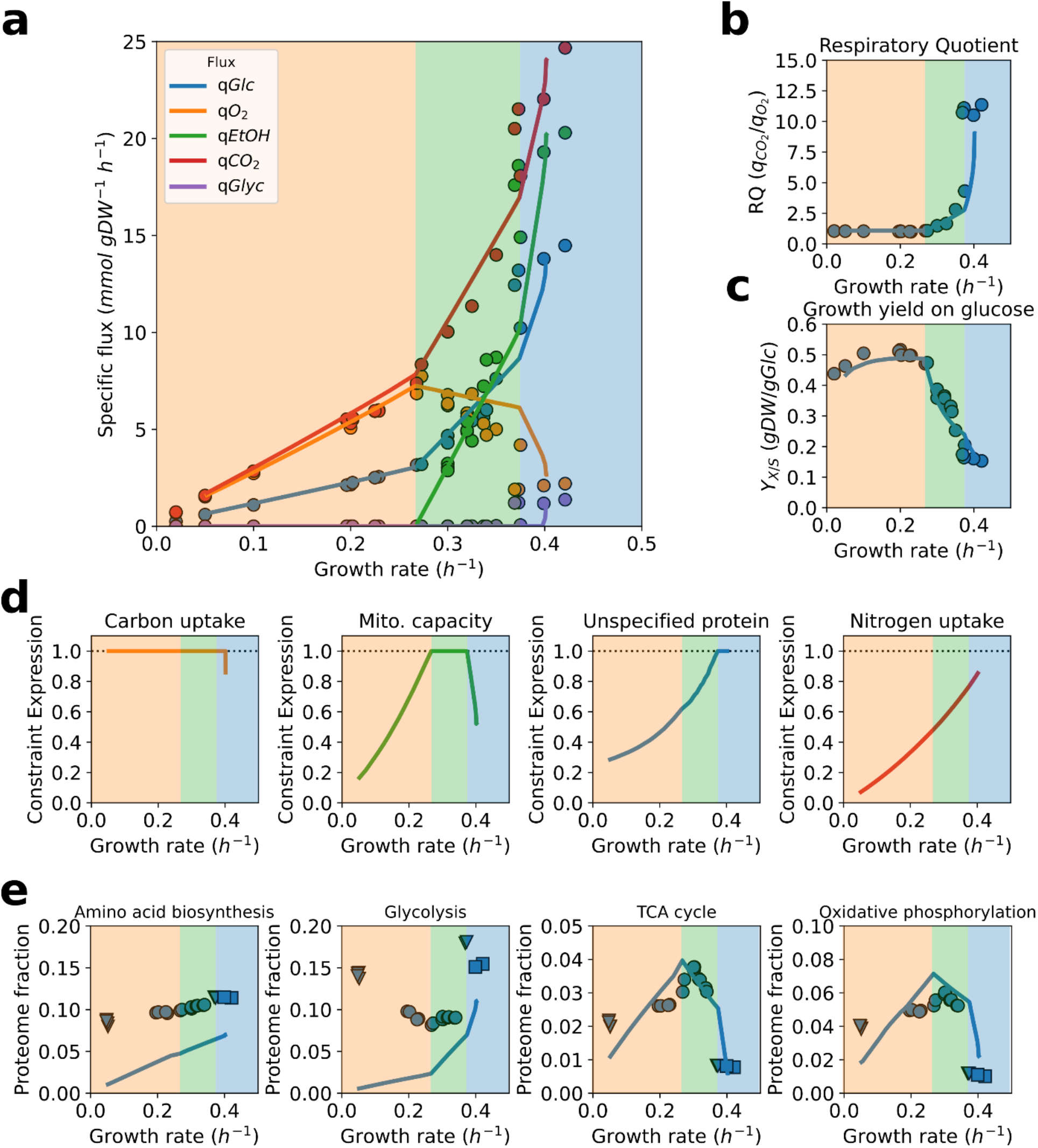
Predicted and measured physiological response of S. cerevisiae CEN.PK as a function of the growth rate in glucose limited aerobic chemostats. **a**. Specific uptake and excretion rates as function of the growth rate. **b**. The respiratory quotient (RQ), the ratio between specific exchange rates of carbon dioxide and oxygen. **c**. Biomass yield on glucose. **d**. Expressions of the model constraints. If the constraint expression equals 1, it is considered active. **e**. Proteome fractions allocated to main pathways in yeast metabolism. The shading in the panels corresponds to the active constraint in Figure 1d. Proteome annotations taken from (Elsemman et al., 2022). Data from glucose-limited chemostat cultures (circles), trehalose- or glucose excess cultures (triangles), and glucose-excess cultures from the control experiment of cycloheximide treatment (squares) from (Elsemman et al., 2022).

To further validate the model predictions, we compared the predicted proteome fractions allocated to specific pathways with quantitative proteomics data (Figure 1e), with predictions similar to these of *pcYeast7*.*6*. The model predicts *minimal* required proteome fractions (assuming that proteins operate at *v*_*max*_). The predicted proteome fractions of TCA cycle and oxidative phosphorylation proteins were in quantitative agreement with the experimental measurements, and captured the qualitative trend for, e.g. amino acid biosynthesis proteins. In contrast to the previous model iteration, the predicted proteome fractions of protein translation-related proteins (translation initiation and elongation factors) were in line with experimental measurements (Supplementary Figure 1). With this we conclude that the improved proteome constrained model of *S. cerevisiae* can give us detailed insight in the physiology of *S. cerevisiae* grown in aerobic, glucose limited chemostat cultures.

### Reduced mitochondrial respiration leads to increased energy costs for growth

With the *pcYeast8* model fully parametrized for glucose-limited aerobic chemostat cultures, we aimed to characterize the physiology of *S. cerevisiae* in energy-perturbed states. Mitochondria are central to energy harvesting in eukaryal cells: these are specialized organelles, which host the TCA cycle proteins and the oxidative phosphorylation system. In yeast, the biosynthesis of mitochondria is mainly regulated by the transcription factor *Hap4* (Maris et al., 2001; Winde and Grivell, 1995). We thus asked whether we can predict changes in yeast physiology due to modulation of *Hap4* expression levels.

We mimicked overexpression and deletion of *Hap4* by adjusting the mitochondrial proteome capacity in glucose-limited chemostats (Figure 2a-b). For the overexpression of *Hap4* (Figure 2a), we needed to increase the mitochondrial capacity to 125% of the wild-type value (Figure 1) to find good agreement in terms of the shifted critical dilution rate *D*_*crit*_ *=* 0.31 *h*^−1^. The subsequent model predicted exchange (uptake and excretion) fluxes across different dilution rates were very close to the observed fluxes.

**Figure 2.**
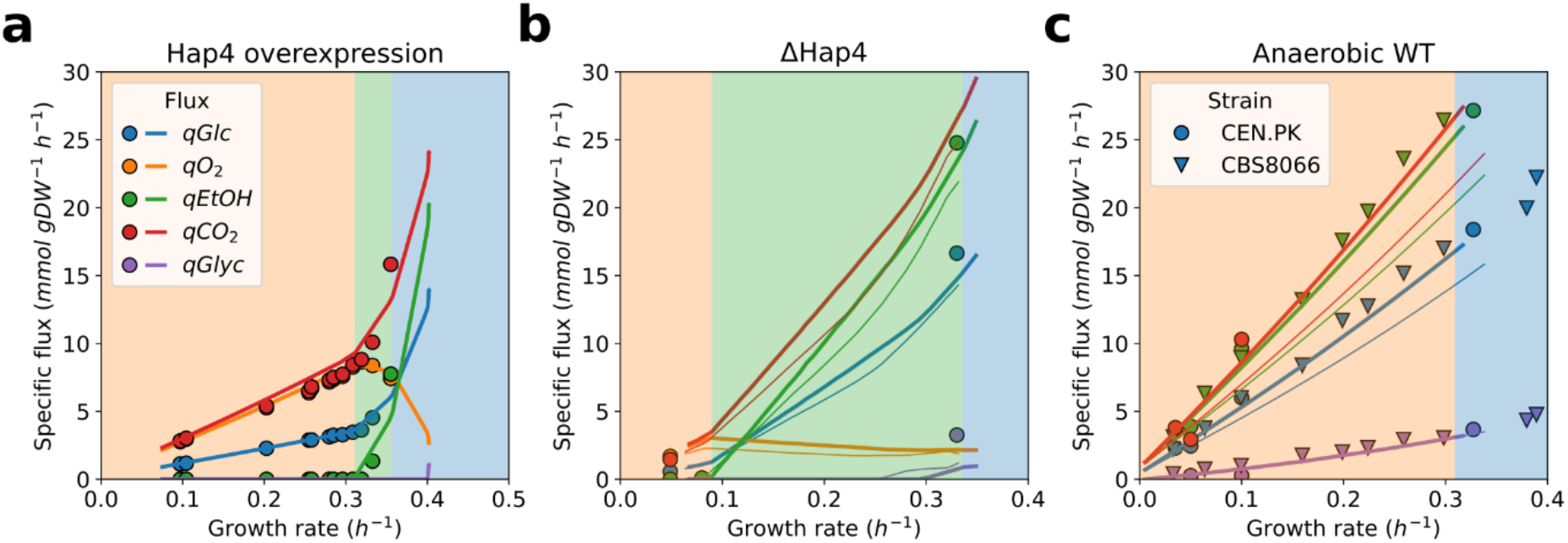
Specific uptake and excretion fluxes as a function of growth rate in glucose-limited chemostat cultures. **a-b** Aerobic growth of the strains with perturbed *Hap4* expression: **a** *Hap4* overexpression strain; **b** *Hap4* deletion mutant (Δ*Hap4*). **c**. Anaerobic growth in glucose-limited chemostats. Points are experimental measurements, lines are model predictions. Shading of the panels corresponds to active proteome constraints at different simulation points as represented in Figure 1. The thicker and thinner lines in **b-c** represent simulations with the growth associated ATP maintenance (GAM) set to 40 and 24 *mmol gDW*^−1^, respectively. Data for **a** from (Maris et al., 2001), for **b** from (Raghevendran et al., 2006), for **c**, two different strains: CEN.PK (circles) from (Björkeroth et al., 2020; Jewett et al., 2013; Tai et al., 2007, 2005), CBS8066 (triangles) from (Nissen et al., 1997).

Less straightforward, however, were the results from an analogous experiment of the *Hap4* deletion mutant (Figure 2b). We had to reduce the mitochondrial proteome capacity to 28% of the WT value to capture the *D*_*crit*_ *=* 0.085 *h*^−1^ of the Δ*Hap4* strain (Raghevendran et al., 2006). Then, we observed that predicted fluxes were lower than measured flux values at the glucose excess conditions (thinner lines in the Figure 2b). This observation called to reassess the ATP maintenance parameters for the Δ*Hap4* strain, the non-growth-associated ATP maintenance (NGAM) and/or the growth-associated ATP maintenance (GAM). Increased NGAM value did not result in more accurate predictions (data not shown), and we found that a substantial increase of GAM to ca. 160 % of the initial (naïve) value (40 vs. 24 *mmol gDW*^−1^) did so (thicker lines of the Figure 2b). With the increased GAM, we then had to readjust the mitochondrial proteome capacity to get to the critical dilution rate and arrived at 32 % of the wild-type proteome capacity. Yet, the ratio of ethanol produced per glucose consumed *q*_*EtOH*_/*q*_*Glc,*_ remained largely unaltered because of the changed GAM value (Figure S3).

Based on our observations, here we conclude that growth with reduced mitochondrial respiration leads to additional energy expenditures, which are currently not captured by the *pcYeast8* model. The improved predictions were achieved by changing global model parameters, in this case, growth-associated ATP maintenance (GAM).

### Additional energetic and proteome investments in anaerobic growth

We wondered if the increased energy costs for growth was specific for Hap4 deletion under glucose excess only or was more generic for loss in respiratory capacity. We therefore used our model to analyze experimental data of anaerobic glucose-limited chemostats (Figure 2c). As with the Δ*Hap4* strain, we tested growth with both the naïve (24) and increased (40 *mmol gDW*^−1^) GAM values. We observed that the same increase in the GAM value, 24 to 40 *mmol gDW*^−1^, was needed to capture the exchange fluxes in anaerobic glucose-limited chemostats (see thicker vs. thinner lines in the Figure 2c). However, we observed that the *in silico* maximal growth rate (of *µ =* 0.38 *h*^−1^) largely exceeded the experimental value (*µ*_*max*_ *=* 0.32 *h*^−1^, (Björkeroth et al., 2020)) for the CEN.PK strain - even with the GAM value of 40 *mmol gDW*^−1^ (Figure S4). We looked at the predicted active constraints for anaerobic growth and identified that to arrive to the maximal growth rate, characteristic to the CEN.PK strain, the minimal level of the “unspecified protein” (UP) had to be increased from the initial value of 0.22 to 0.32 *g UP* (*g protein)*^−1^.

The UP is an artificial protein of average composition and length, and its expression represents all proteins that do not contribute actively to biomass production (i.e. metabolically inactive proteins). The minimal UP mass fraction is, therefore, a proxy for total cytosolic proteome capacity, which becomes an active constraint when the UP fraction in the proteome reaches this preset minimal value. At that point, each protein in the cytosol contributes directly to growth, and investment in one protein has to come at the expense of another. Unlike the increased energy demand, the increase in the UP value is only relevant for anaerobic growth, and suggests lower “usable” proteome capacity even at glucose excess.

Overall, the combination of the model parameters we had to change *explicitly* to capture the anaerobic physiology of *S. cerevisiae* suggests the presence of additional energy and proteomic demands associated with growth when no or a very limited amount of oxygen can be used for oxidative phosphorylation.

### Predicting growth of *S. cerevisiae* in rich media

The growth-associated ATP maintenance and the minimal UP level that we needed to adjust to capture anaerobic growth, are global parameters, estimated from available data but under uncertainty and assumption. We thus wanted to test whether the parameter values we adopted lead to a reasonable prediction of growth under different growth conditions, for which we studied a nutritional upshift.

A major contributor to the energy and proteome costs for growth in minimal, chemically-defined, media is the biosynthesis of amino acids. Later polymerized into proteins, proteins are the most abundant constituent of yeast dry biomass (between 35 and 50 % of the total dry mass, (Canelas et al., 2011)) and protein turnover is a major consumer of ATP in the cell (Lahtvee et al., 2017). It is thus unsurprising that, compared to minimal media, *S. cerevisiae* exhibits a ca. 20 % higher specific growth rate in rich media, such as YPD or SC (Metzl-Raz et al., 2017).

We analyzed the experiment performed by (Björkeroth et al., 2020): growing *S. cerevisiae* in either minimal Verduyn medium, or medium supplemented with amino acids (rich medium), in both aerobic and anaerobic conditions. We estimated the transporter capacity and uptake of amino acids for the rich medium as detailed in Methods, and predicted batch growth in these four conditions (Figure 3).

**Figure 3.**
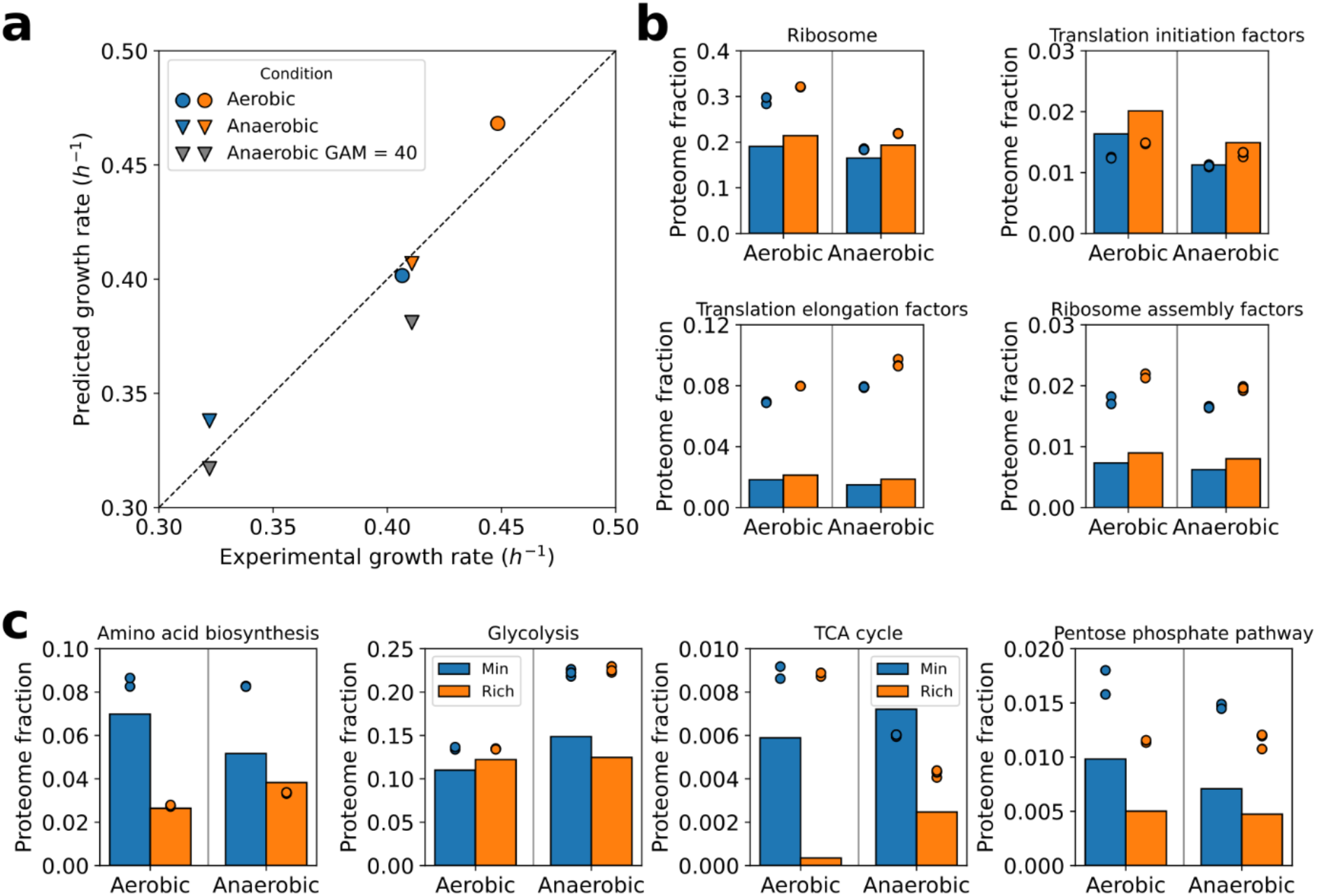
Overview of the predicted growth rates and proteome composition in batch cultures with glucose as the main carbon source. **a**. Predicted batch growth rates for minimal and amino acid-rich media. **b-c** Protein abundance in different conditions (mass fractions *g* (*g protein)*^−1^): **b**. translation-related proteins, **c**. clusters of energy- and amino acid metabolism. Blue and orange points/bars represent minimal and rich media, respectively. Points in **b-c** are experimental measurements, bars are model predictions. Proteome annotations taken from (Elsemman et al., 2022). Data from (Björkeroth et al., 2020).

We first simulated growth using the GAM values as determined previously: 24 *mmol gDW*^−1^ for aerobic and 40 *mmol gDW*^−1^ for anaerobic growth. The predicted batch growth rates were similar to the experimentally measured values (Figure 3a) for 3 out of 4 conditions, with the exception of the underpredicted anaerobic growth rate on the rich medium. For this condition, we reversed the GAM value to 24 *mmol gDW*^−1^ and observed an increased predicted growth rate which was more consistent to the experimental measurements. Consequently, we have adopted the GAM value of 40 *mmol gDW*^−1^ for anaerobic growth on minimal Verduyn medium only.

In the proteome predictions, we have observed a growth rate-dependent increase in the proteome fraction, allocated to ribosomes (Figure 3b). The predictions were in agreement with previous reports (Elsemman et al., 2022; Metzl-Raz et al., 2017), however, the absolute ribosomal proteome fraction in aerobic conditions in the dataset of (Björkeroth et al., 2020) was considerably higher as reported of (Elsemman et al., 2022; Metzl-Raz et al., 2017). Nonetheless, the relative upshift of ribosomes in amino acid-supplemented media was predicted correctly (Figure S5).

Other translation-related non-ribosomal proteins (translation factors *etc*.) exhibit a growth rate-dependent increase as well (Figure 3b, similar to the increase due to glucose availability, Figure S1). A notable exception between data and predictions was the reported fraction of translation elongation factors, which was ca. 3-fold higher than both the model predictions (Figure 3b), and our previously published data from aerobic batch cultures in minimal media (Figure S1). We do not have explanations for the high abundance of translation elongation factors and ribosomes in the particular dataset of (Björkeroth et al., 2020), and we speculate that this could be perhaps dictated by technical bias.

In glucose batch conditions, where the cytosolic proteome constraint is active, the increase in the proteome spent on protein translation has to be accompanied by a decrease in other proteome clusters. The major, and the most obvious change for growth on rich medium was the ca. 2-fold decrease in proteins associated with amino acid biosynthesis, as well as TCA cycle proteins (Figure 3c). Some other proteins relevant to the biosynthesis of amino acids, such as pentose phosphate pathway, were also less expressed in rich media. Contrary to that, as expected, the abundance of glycolysis proteins did not significantly vary because of amino acid supplementation.

To sum up, here we challenged the *pcYeast8* model to rich, amino acid-supplemented media in both aerobic and anaerobic conditions. The current estimates of the GAM and minimal UP values for growth in both aerobic- and anaerobic conditions allowed us to successfully predict physiology and proteomes for growth in rich media, using an independent dataset. The predicted additional energy investment, needed for anaerobic growth on a minimal medium, was alleviated by supplementation of amino acids to the medium. Together with the previous observations in the aerobic Δ*Hap4* cells, these results suggest that the additional costs of growth in anaerobic conditions might stem from the requirement of functional mitochondria for the *de novo* biosynthesis of amino acids.

### Redox shuttling via acetaldehyde-ethanol shuttle involves additional costs

Since the GAM value is a global parameter with no mechanistic role in the *pcYeast8* model, we next wanted to explore possible origins of increased cost of generating new biomass under anaerobic conditions – new amino acids for proteins, to be specific. One specific challenge in anaerobic/respiratory-deficient growth is maintaining redox balance in cell compartments. Synthesis of cellular components creates a surplus of reduced electron carriers, such as NAD(P)H, in both cytosol and mitochondria. Aerobically, cytosolic redox equivalents are shuttled into mitochondria and fed into the electron transport chain. Additionally, in fast aerobic growth (Figure 1a), some of the NADH is reoxidized in the glycerol shunt (production of glycerol from dihydroxyacetone phosphate).

Yet in anaerobic conditions, the only available option to resolve redox imbalances is via glycerol production in the cytosol (Figure 2c, (Nissen et al., 1997)), and shuttling of redox equivalents out of mitochondria becomes essential for growth. There are three main redox shuttles connecting cytosolic and mitochondrial NAD^+^/NADH pools in *S. cerevisiae* (Bakker et al., 2001); two of them are active in aerobic conditions, and the acetaldehyde-ethanol shuttle (operated by the mitochondrial alcohol dehydrogenase *Adh3*) is also present anaerobically (Bakker et al., 2000) (Figure 4b). Notably, the *pcYeast8* model predicted no additional costs linked to the shift from oxygen-dependent shuttles to acetaldehyde-ethanol shuttle (Figure 2c, thin lines). So we asked how the predicted physiology changes depending on the shuttle used.

**Figure 4.**
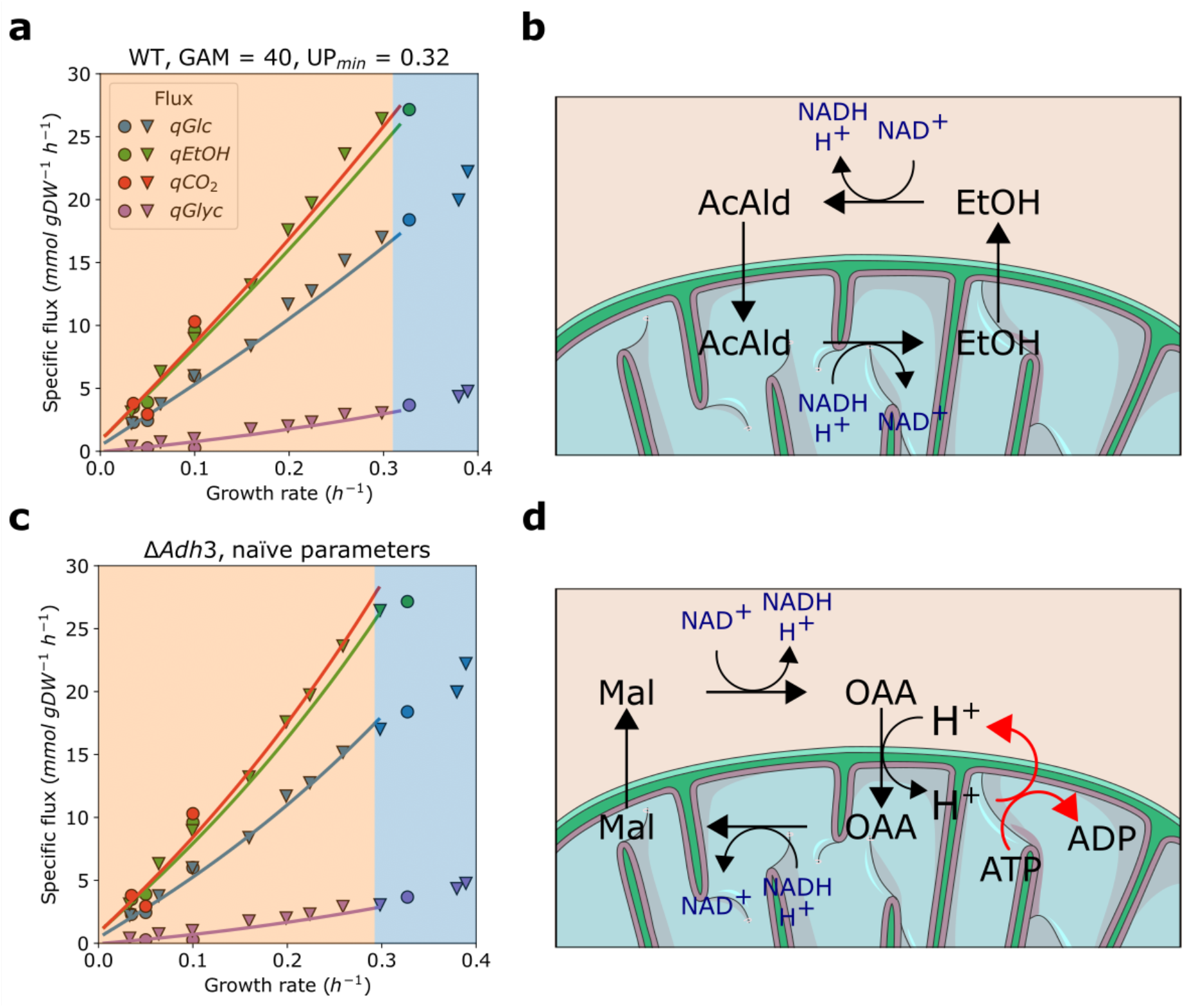
Specific uptake and excretion fluxes as a function of growth rate in anaerobic glucose-limited chemostat cultures with different redox shuttles in mitochondria. **a-b** Physiology of wild-type cultures with the parameters changed (see Fig. 2c). In the absence of oxygen, model predicts the acetaldehyde-ethanol shuttle (**b**) to be active. **c-d** Physiology of the deletion mutant of the mitochondrial alcohol dehydrogenase *Adh3*, where oxaloacetate-malate shuttle (**d**) is predicted to shuttle NADH from mitochondria to cytosol. Points in **a** and **c** are experimental measurements (see Fig. 2c for legend and sources), lines are model predictions. The following parameter values were used for a and c, respectively: GAM, 40 and 24 *mmol gDW*^−1^; UP, 0.32 and 0.22 *g UP* (*g protein)*^−1^.

As introduced previously, for cells using the acetaldehyde-ethanol shuttle, we needed to set additional energy and proteome costs order to tailor the flux predictions to experimental observations (Figure 2c, reproduced in Figure 4a). The activity of the shuttle does not result in transfer of additional metabolites (such as protons) across the mitochondrial membrane (Figure 4b). However, it is known that high concentration of alcohols results in significant increase in proton permeability of membranes (Leão and Van Uden, 1984). Yet the magnitude of the ethanol-induced permeability (i.e. stoichiometry of protons crossing the membrane per cycle of acetaldehyde-ethanol shuttle) is not known (and likely hard to measure experimentally). We have estimated that import of 3 protons/cycle to mitochondria could eventually explain the increase in energy demand (data not shown), but we believe such a high ratio is unlikely and thus reject this being the only factor linked to these costs. Moreover, the specific excretion rate of ethanol between aerobic vs. anaerobic glucose batch (Figure 1a vs. Figure 2c) varies within 30% range, and so permeabilization would also lead to additional energetic costs in aerobic batch cultures, although we have no supporting data (e.g. need to change model parameters) to confirm this.

Since operating the acetaldehyde-ethanol shuttle seems to be “too cheap” in the model, are there any alternatives? The cells depleted of *Adh3* use another oxygen-independent option for redox balancing, the malate-oxaloacetate shuttle ((Bakker et al., 2001), Figure 4d). In fact, it is an energy-costly option since it involves oxaloacetate-proton symport to mitochondria: the imported protons must be pumped out back to the cytosol by the mitochondrial ATPase at the cost of 1 ATP per proton. Surprisingly, use of the malate-oxaloacetate shuttle does not require any changes to the global parameters (GAM and minimal UP value) to predict both the fluxes and the maximal growth rate in glucose excess within correct range (Figure 4b).

Existing experimental data, however, highlights the role of the acetaldehyde-ethanol shuttle in anaerobically growing cells ((Bakker et al., 2000) and discussed below). Based on this, we cannot assume that the Δ*Adh3* flux profile (use of an alternative shuttle with extra costs) corresponds to the *actual* wild-type behavior. There might be multiple explanations and assumptions we could undertake (see Discussion). However, in sake of consistency, in the current study we will further consider the case with the acetaldehyde-ethanol shuttle present and model parameters adjusted as *the* wild-type profile.

### Effective redox shuttling in and out of mitochondria is required for fast anaerobic growth

We have previously observed that redox shuttling from mitochondria to cytosol is key for anaerobic growth, where mitochondria serve mainly biosynthetic functions. Moreover, some of the biosynthetic processes in mitochondria use NADPH, the majority of which is supplied by the malic enzyme *Mae1* in anaerobically-growing cells (Boles et al., 1998). We thus wanted to stratify the importance of these two systems in anaerobic, glucose-limited growth, and analyzed the predicted impact of deletion of *Mae1* and *Adh3* on the physiology of the anaerobic-growing *S. cerevisiae* (Figure 5).

**Figure 5.**
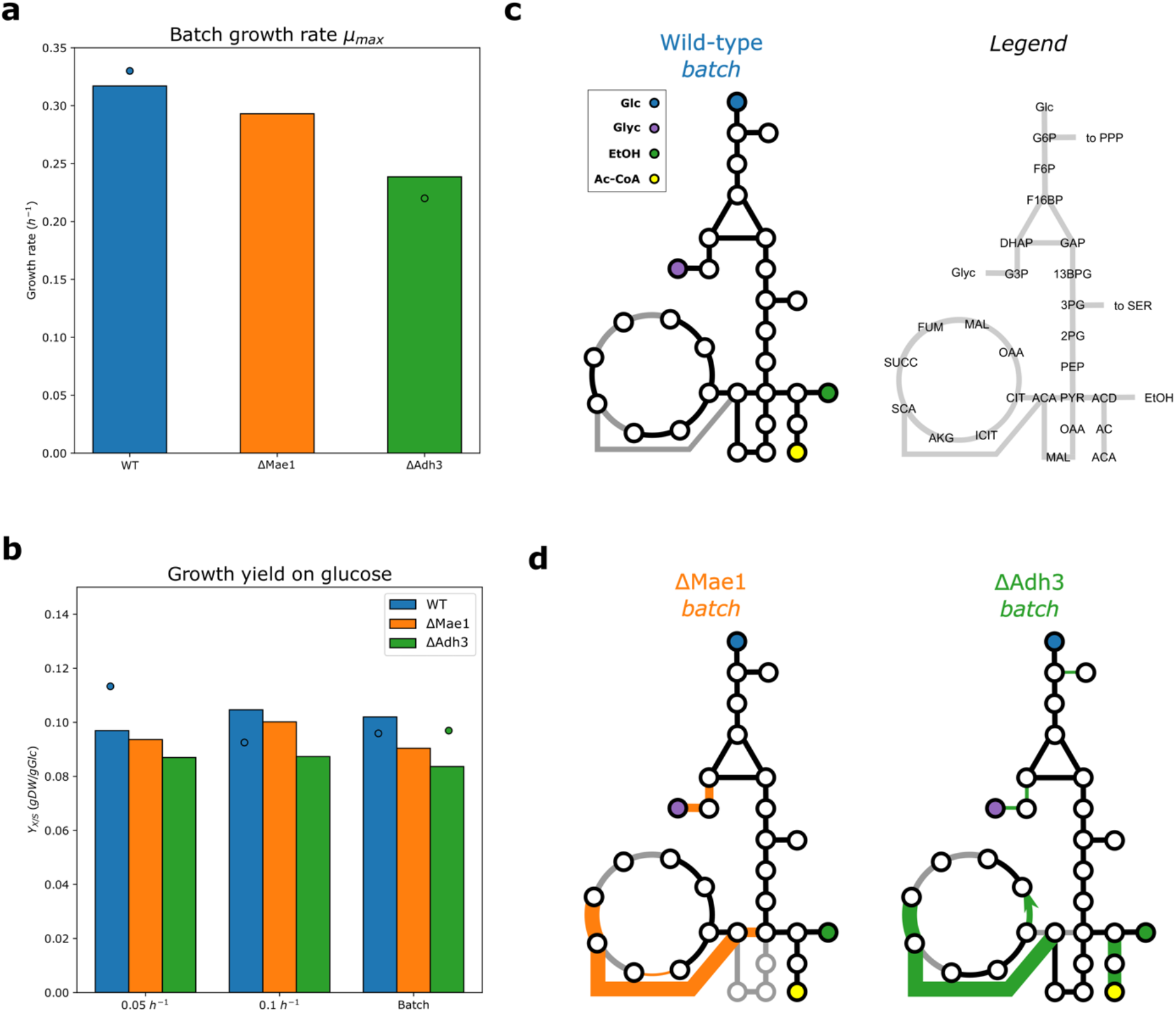
Physiological parameters and predicted intracellular fluxes for anaerobically growing deletion strains and wild-type strain in nitrogen-limited chemostats. **a-c** Deletion strains of the malic enzyme Δ*Mae1* and mitochondrial redox shuttle Δ*Adh3*. **a** Predicted maximal growth rate in anaerobic batch cultures. **b** Predicted yield on glucose in anaerobic glucose-limited chemostats at *D =* 0.05 *h*^−1^, *D =* 0.1 *h*^−1^ and batch cultures. **c-d** Comparison of the predicted flux profiles in the central carbon metabolism. Line width in **d** represent relative flux value (maximal thickness 2.5-fold of the wild-type value), with the wild-type batch (**c**) as reference. Highlighted in orange or green are fluxes which differ by at least 30 % from the wild-type value, and grey lines indicate inactive fluxes. Data for **a-b** from (Bakker et al., 2000; Björkeroth et al., 2020; Boles et al., 1998).

It should be noted that in the *pcYeast8* model, the glycerol 3-phosphate shuttle (*Gpd2*) is respiration-independent and has to be knocked-out *in silico* together with *Adh3* for predicting the phenotype of experimental Δ*Adh3* mutants (model predictions further called just Δ*Adh3* instead of Δ*Adh3*Δ*Gpd2*). Both *Mae1* and *Adh3* deletion were reported to have very little impact on the physiology of glucose-limited chemostats at low dilution rates (Bakker et al., 2000; Boles et al., 1998), and this claim is supported by our model predictions (Figure S6). Thus we focused on the batch growth of the Δ*Mae1* and Δ*Adh3* mutants instead.

First, we looked at the predicted maximal growth rate (Figure 5a) and yield on glucose (Figure 5b) in anaerobic cultures of Δ*Mae1* and Δ*Adh3* mutants. The batch growth rate for the Δ*Mae1* mutant was slightly reduced compared to the wild-type strain, in agreement with experimentally reported “little-to-no impact on the physiology by the *Mae1* deletion” (Boles et al., 1998). Meanwhile, the Δ*Adh3* strain showed a ca. 1/3 reduction of the batch growth rate, consistent with experimental reports (Bakker et al., 2000). The predicted growth yield on glucose (Figure 5b), however, was lower for the Δ*Adh3*, compared to the experimentally determined value, which was comparable to the wild-type yield.

To understand the differences in growth rates and yields, we also looked at the predicted intracellular fluxes (Figure 5c, 5d), with a focus to the central carbon metabolism. The main difference between the predicted flux distributions of the wild-type and mutant strains is that the mutant strains generate a lot more (ca. 20*×* the WT flux) mitochondrial ATP using one of the TCA cycle reactions, succinyl-CoA ligase (succinyl-CoA + ADP + Pi succinate + CoA + ATP). For running this ATP generation route, the coenzyme A moiety is transferred from the acetyl-CoA to succinate (succinate + acetyl-CoA succinyl-CoA+ acetate), which is then regenerated in the succinyl-CoA reaction. What differs between the mitochondrial ATP generation in Δ*Mae1* and Δ*Adh3* strains is the predicted source of mitochondrial acetyl-CoA: the Δ*Mae1* mutant supplies the acetyl-CoA via decarboxylation in the mitochondria by pyruvate dehydrogenase complex, while Δ*Adh3* uses cytosolic acetyl-CoA, shuttled to mitochondria using the carnitine shuttle.

Normally, acetyl-CoA is produced in the cytosol in low amounts for the biosynthesis of fatty acids. In the Δ*Adh3* strain, cytosolic acetyl-CoA production is increased multiple-fold, even though this is an energy-spilling route: for one ATP generated in mitochondria, 2 equivalents of ATP are consumed in cytosol (acetate + CoA + ATP *→* acetyl-CoA + AMP + PPi). However, the reduced NADH (gained through the aldehyde dehydrogenase reaction, acetaldehyde + NAD^+^ *→* acetate + NADH + H^+^) remains in the cytosol, and a proton (because respective molecule of pyruvate is not imported into mitochondria) is not imported to mitochondria. The amount of excess NADH produced scales with the growth rate, as indicated by decreasing NAD/NADH ratio as the growth rate increases (Canelas et al., 2011), and the high additional demand for energy to resolve the redox imbalances could explain the lower batch growth rate of the Δ*Adh3* strain. This, however, stands true only to growth on minimal medium: the predicted batch growth rate of the Δ*Adh3* mutant on a rich medium was only mildly lower than the predicted wild-type rate (Figure S7).

## Discussion

In this study, we presented the updated proteome-constrained model of the budding yeast *Saccharomyces cerevisiae, pcYeast8* (Figure 1). We have built a proteome-constrained model on the basis of the genome-scale model (GEM) *Yeast8* (Lu et al., 2019), inspired by our recently published model *pcYeast7*.*6* (Elsemman et al., 2022). We manually curated some of the model reactions (Supplementary Notes) which allowed correct prediction of some growth strategies, e.g., glycerol excretion in aerobic, glucose-excess conditions (Figure 1), which were not properly captured by the *pcYeast7*.*6* yet.

The focus of our study was to identify condition-dependent patterns of yeast physiology, with extra attention to the energy expenditures for growth. We first modulated the respiratory capacity of *S. cerevisiae*, and tested scenarios of increased, restricted, or completely blocked respiration (Figure 2). Here we identified two model parameters that must be tweaked in order to capture the uptake and excretion fluxes in respiration-deficient (Δ*Hap4*) mutants, or anaerobically grown cells (Figure 2b, 2c): the growth-associated ATP maintenance (GAM) and the minimal fraction of the proteome, to be occupied by the so-called unspecified protein (UP). In fact, despite the size and extent of the *pcYeast8* model, only a handful of global model parameters can be effectively used for fitting: both in the previous model iteration (Elsemman et al., 2022), and in our current study, we addressed three global parameters: the uptake of carbon source (see Methods), the minimal UP fraction in proteome, and growth-associated ATP maintenance. We have validated the choice of the parameter values for the minimal medium in an experiment of growth in rich medium (Figure 3), where we successfully predicted the batch growth rates and proteome compositions across conditions.

Both aerobic Δ*Hap4*, and anaerobic *S. cerevisiae* exhibit reduced mitochondrial content (Figure S2); however, mitochondria remain important – or even essential – for some growth processes. The significant reduction in the biomass yield per mol ATP produced (*Y*_*ATP*_), comparing anaerobic glucose-limited chemostats vs. aerobic batch cultures was already observed by (Verduyn et al., 1990b), especially at higher dilution rates in anaerobic chemostats. The additional energy expenses for growth on minimal medium, however, are very high (an increase in the GAM value by 16 *mmol gDW*^−1^, or a 60% increase of the naïve value. Our model predicted that supplementation of amino acids rescued the extra energy requirement in anaerobically-grown *S. cerevisiae* (Figure 3a). Inhibited respiration in fission yeast *Schizosaccharomyces pombe* resulted in reduced proliferation and decreased levels of 4 amino acids (Arg, Lys, Glu, Gln) (Malecki et al., 2020); supplementation of Arg to the medium rescued the growth reduction. Our modeling results are in line with these observations, since the increase of GAM from 24 to 40 *mmol gDW*^−1^ was not needed for correctly predicting the physiology of anaerobically-grown *S. cerevisiae* in rich medium (Figure 3a).

A combination of several factors, e.g., increased futile cycling or decoupling of energy generation from biomass formation, were proposed to contribute to additional costs, usually without clear mechanistic insight. Some studies highlighted the function of mitochondria in biosynthesis (Visser et al., 1994) and lipid metabolism (Reiner et al., 2006) and suggested that maintaining functional mitochondria in anaerobic conditions is an energy-costly process (Drgoň et al., 1991; Visser et al., 1994). Moreover, recent reports allow us to speculate that maintenance of proton-motive force across cell membranes could be one of the factors leading to extra costs in anaerobic growth ((Malina et al., 2021; Terradot et al., 2021),

T. Pilizota, personal communication). Aerobically, the proton-motive force is generated by the electron transport chain in the mitochondrial inner membrane, and some of it is used for, e.g. import of metabolites and proteins into mitochondria. In anaerobic settings, not only the proton-motive force is not maintained by the electron transport chain, but also can be eradicated due to increased permeability of membranes due to increased intracellular ethanol concentration (Leão and Van Uden, 1984).

We speculated that these energy costs might be associated with impaired redox shuttling in mitochondria. Redox equivalents can be shuttled using two (acetaldehyde-ethanol and malate-oxaloacetate) shuttles in anaerobic growth. Yet the simulations of the preferred (acetaldehyde-ethanol) shuttle were contradictory to experimental data as we had to change global model parameters to capture experimental data (Figure 2c, Figure 4a). Meanwhile, use of alternative redox shuttle led into predictions consistent with experimental measurements (Figure 4c). This contradiction could be linked to different putative roles of the preferred redox shuttle, e.g., potential moonlighting roles of *Adh3* enzyme, or simultaneous shuttling through both acetaldehyde-ethanol and malate-oxaloacetate shuttles – all of which are still open for further investigation.

We did scenario testing with respect to the maintenance of redox balance in anaerobic mitochondria (Figure 5). We have simulated batch growth of a mitochondrial redox shuttle-deficient yeast and obtained predictions of batch growth rate in agreement with experimental data (Figure 5a, (Bakker et al., 2000)). We suggest that the increased cytosolic decarboxylation of pyruvate at the expense of ATP (Figure 5c, 5d) can sustain biosynthesis and partially alleviate the redox imbalance in mitochondria, but not at high growth rates, when the glycolytic flux is very high. Predicted batch growth rate of Δ*Adh3* mutant in rich medium was very comparable to the wild-type (Figure S7), and, again, pointed out to the reduction of growth being related to amino acid biosynthesis. However, the exact mechanisms of the energy-consuming processes, characteristic to low- or non-respiring mitochondria, however, remain to be elucidated.

The additional cost of proteome maintenance in anaerobic conditions (Figure 2c) is also an important consideration. In our model, we express the maximal capacity of “useful” (=involved in biomass production) proteome as the minimal proteome fraction of the “dummy” protein, UP. We had to increase the minimal fraction of UP in order to observe a batch growth rate, corresponding to the experimentally observed growth of *S. cerevisiae* CEN.PK strain.

We speculate the additional proteome space is allocated to anticipatory protein expression, mainly, proteins needed for growth in the presence of oxygen. For instance, a substantial overexpression of the oxidative phosphorylation enzymes was observed in microaerobic conditions (Rintala et al., 2009). In the lab settings, unsaturated fatty acid- or ergosterol are supplemented to anaerobic cultures of *S. cerevisiae* to override the need for oxygen-requiring biosynthesis steps. Anticipation of presence of oxygen might also explain why *S. cerevisiae* can grow anaerobically, albeit slowly, without unsaturated fatty acid- or ergosterol supplementation, which is a common technique in laboratory cultivation (Dekker et al., 2019). In general, cells show anticipatory expression of metabolic enzymes when the selection pressure for the *transient* growth rate is absent: we have recently suggested a similar phenomenon for glycolytic enzymes for aerobic glucose-limited cultures (Grigaitis and Teusink, 2022). Also, glycolytic enzymes show similar levels in both minimal and rich media despite the differences in glycolytic flux (Björkeroth et al., 2020).

To conclude, in this study we have used the proteome-constrained model of yeast metabolism to explore the metabolic strategies of *S. cerevisiae* across different growth conditions. We have identified condition- and mutant-specific costs of growth, which we associated with impaired mitochondrial respiration and redox shuttling. Some of these costs have direct mechanistic explanations, coming out of the predictions of the *pcYeast8* model; for some, the exact underpinnings are still open for future studies. In the end, we hope that our modeling work will open some new avenues for the research of yeast metabolic- and resource allocation strategies.

## Methods

### Kinetics and proteome data for the *pcYeast8*

The detailed description of reconstruction of the proteome-constrained model of *S. cerevisiae* is provided in the Supplementary Notes. 5’-UTR sequences and proteome annotations (composition of macromolecular complexes, Gene Ontology terms etc.) were collected from Saccharomyces Genome Database (SGD, (Cherry et al., 2012)).

We used the reference proteome of *S. cerevisiae* from UniProt (The UniProt Consortium et al., 2021). The kinetic data (enzyme turnover values) were collected from the BRENDA database (Chang et al., 2021). For every enzymatic complex with an Enzyme Commission (EC) number, we queried the BRENDA database for a value from the wild-type enzymes. When available, values from *S. cerevisiae* were preferentially selected. Otherwise, the highest value for a wild-type enzyme in mesophilic (and close to growth conditions of *S. cerevisiae*) conditions was taken. When no *k*_*cat*_ value was available, we assumed *k*_*cat*_ *=* 71*s*^−1^ as a default value (Nilsson and Nielsen, 2016). If the experimentally determined *k*_*cat*_ value was lower than 1*s*^−1^, we set this value.

### Model simulations

Unless stated otherwise, all simulated media were Verduyn minimal medium (Verduyn et al., 1992). Growth in a specific medium was modeled by closing all reverse exchange (“uptake”) reactions, except for those nutrients which are present in the medium (the upper bound of the exchange reaction 0). In the case of anaerobic growth, minute amounts of ergosta-5,7,22,24(28)-tetraen-3beta-ol, oleate, and palmitoleate were also provided for the biosynthesis of unsaturated fatty acids (mimicking the addition of ergosterol and Tween 80 in respective experiments).

Glucose limitation was simulated by altering the saturation of respective plasma membrane transporters, 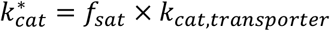. To prevent the plasma membrane from being fully filled with transporters, the available membrane area which these transporters was set to a limited fraction of the total plasma membrane surface area (Table 1).The definition for the rich medium (Verduyn + amino acids) was determined on the basis of data from batch cultures (Björkeroth et al., 2020). There, the time evolution of biomass concentration in the growth medium (*gDWL*^−1^) and amino acid abundance (*µM*) was reported. We computed the slope of depletion of every amino acid in the media (=uptake by cells) and multiplied by the specific growth rate *µ* to determine the upper flux bound for each amino acid quantified. When the slope was nonnegative (no uptake in essence), we set the upper flux to zero.

**Table 1.** Plasma membrane transporter area for different growth conditions.

| Condition | Carbon transporters, % | Nitrogen transporters, % |
| --- | --- | --- |
| Aerobic, minimal medium | 11.0 | 4.5 |
| Aerobic, rich medium | 13.0 | 5.5 |
| Anaerobic | 13.0 | 4.5 |

### Software

The *pcYeast8* model was simulated using the CBMPy package (version 0.8.1) (Olivier et al., 2021) in a Python 3.6 environment with the IBM ILOG CPLEX Optimization Studio (version 12.10.0) and SoPlex (version 5.0.2) (Gleixner et al., 2018) as the low- and high-precision LP solver, respectively. In SoPlex, the primal (−*f* flag) and dual (−*o* flag) feasibility tolerance was set to 10^−16^. R (version 3.6.3) and Python (version 3.10) were used for further data analysis and derivation of relations between the cell size and biomass composition as a function of the growth rate.

## Data Availability

The *pcYeast8* model, and the materials to generate *pcYeast8* model, together with information, required to generate the figures of this manuscript, are available on Zenodo [10.5281/zenodo.7322850] (Grigaitis et al., 2022).

## Funding

PG and BT acknowledge the funding by Marie Skłodowska-Curie Actions ITN “SynCrop” (grant agreement No 764591). We thank SURFsara for the HPC resources through access to the Lisa Compute Cluster.

## Acknowledgements

We thank Frank J. Bruggeman and Jack Pronk for discussions, and Ursula Kummer for her comments on the draft version of the manuscript.

## Competing Interests Statement

None declared.

## CRediT Author Role Statement

Pranas Grigaitis: conceptualization, methodology, software, validation, formal analysis, investigation, data curation, writing – original draft, writing – review & editing, visualization, supervision; Samira L. van den Bogaard: methodology, validation, formal analysis, investigation, writing – review & editing; Bas Teusink: supervision, project administration, funding acquisition, writing – review & editing.

## Notes

### Competing Interest Statement

The authors have declared no competing interest.

