## Supplementary Figures for "Elevated energy costs of biomass production in mitochondrial-respiration deficient *Saccharomyces cerevisiae*"

This document includes:

**Supplementary Figures 1-7**

Separate Supplementary files:

**Supplementary Notes** (.pdf)

### Supplementary Figures

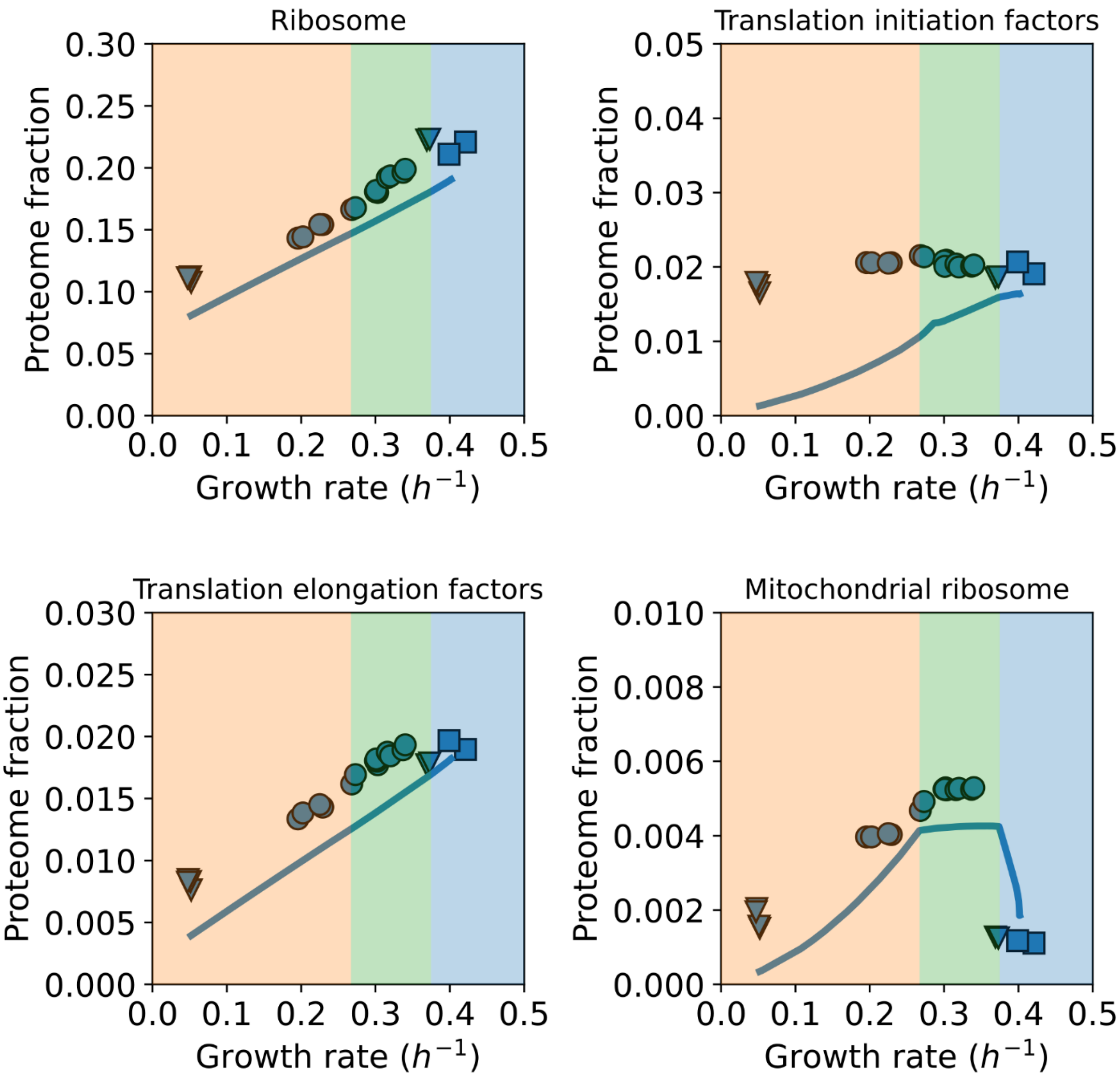


**Supplementary Figure 1. Predicted proteome abundance of translation-related proteins as a function of growth rate in glucose-limited chemostats.** Points are experimental measurements, lines are model predictions of mass fractions in $(g(gprotein)^{-1})$. Shading of the panels corresponds to active proteome constraints at different simulation points as represented in Main Text Figure 1. Proteome annotations taken from (Elsemman et al., 2022). Data from glucose-limited chemostat cultures (circles), trehalose- or glucose excess cultures (triangles), and glucose-excess cultures from the control experiment of cycloheximide treatment (squares) from (Elsemman et al., 2022).

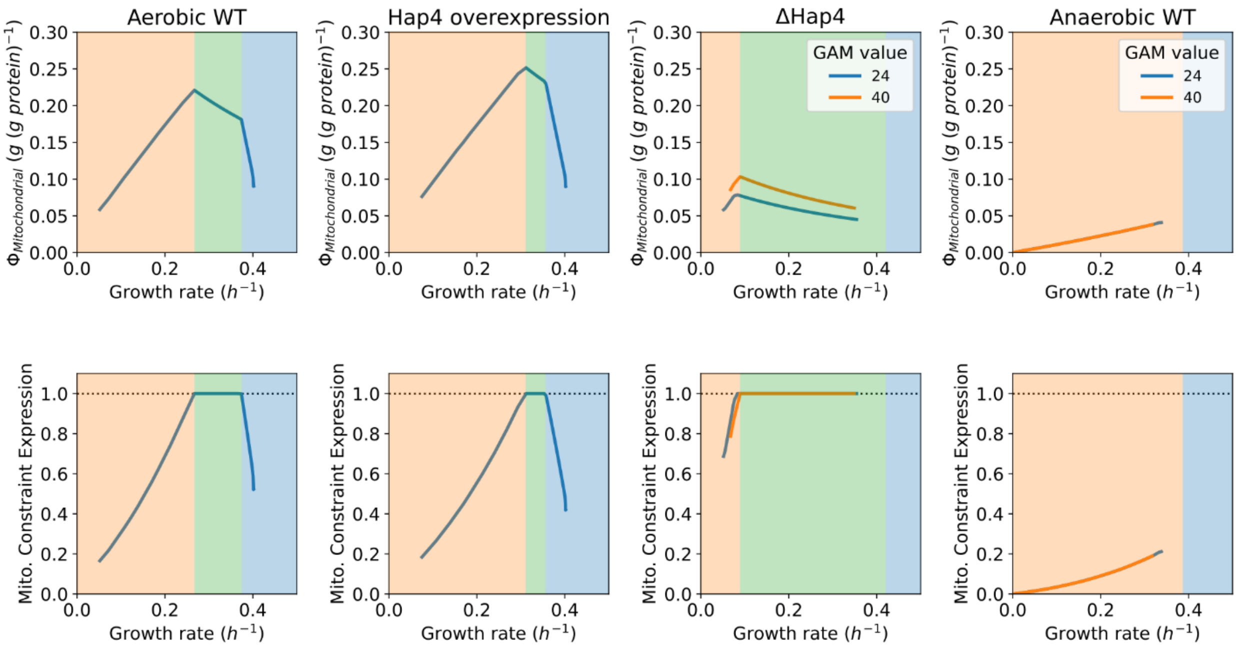


**Supplementary Figure 2. Predicted mitochondrial proteome mass fractions (top row) and mitochondrial capacity constraint expressions (bottom row) in glucose-limited chemostats.** The mitochondrial proteome mass fraction $\Phi_{Mitochondrial}$ is expressed in $(gmprotein(gprotein)^{-1})$. Different colors of the lines in the panels for aerobic $\Delta Hap4$ mutant and anaerobic wild-type strain represent predictions with different growth-associated ATP maintenance (GAM) values (in $mmolgDW^{-1}$). Shading of the panels corresponds to active proteome constraints at different simulation points as represented in Main Text Figures 1 and 2.

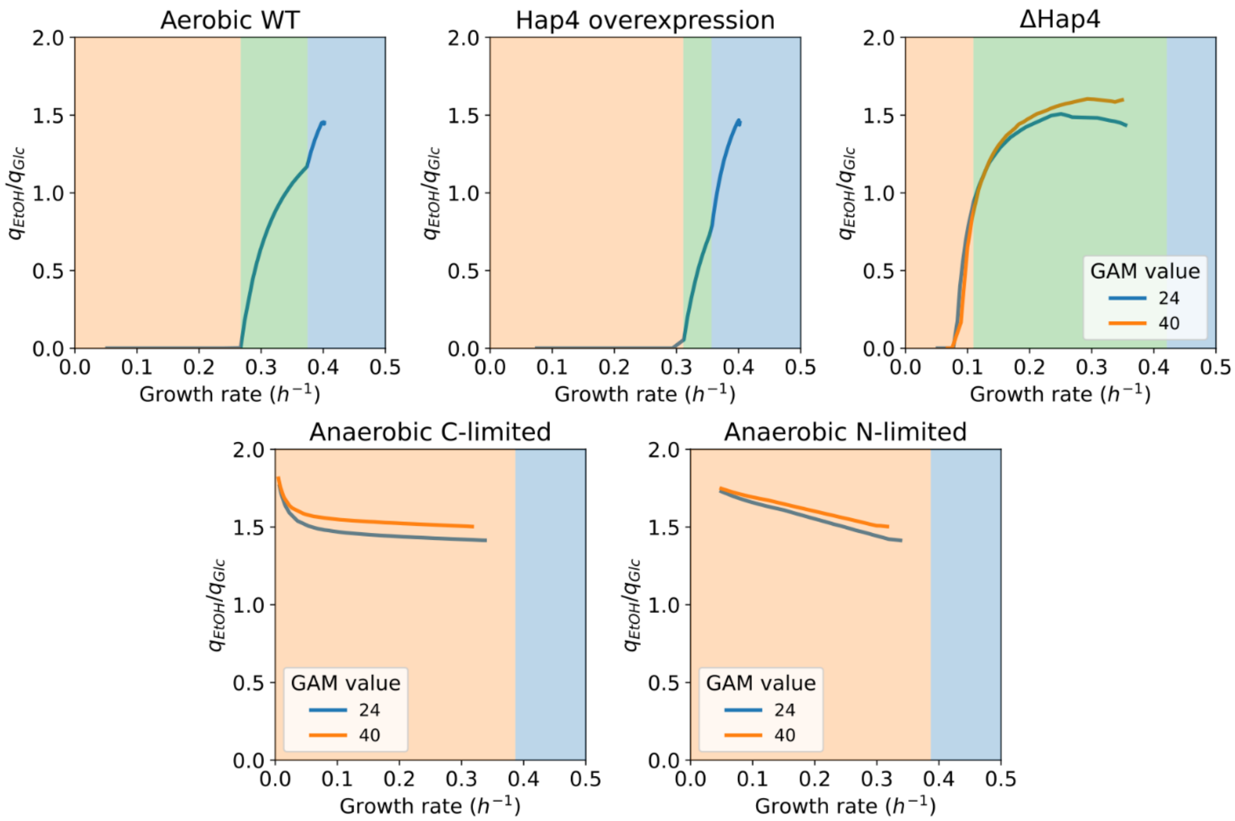


**Supplementary Figure 3. The predicted ratio of excreted glucose vs. glucose consumed as a function of growth rate in glucose- and ammonium-limited chemostats.** Different colors of the lines in the panels for aerobic $\Delta Hap4$ mutant and anaerobic wild-type strain represent predictions with different growth-associated ATP maintenance (GAM) values (in $mmolgDW^{-1}$). Shading of the panels corresponds to active proteome constraints at different simulation points as represented in Main Text Figures 1 and 2.

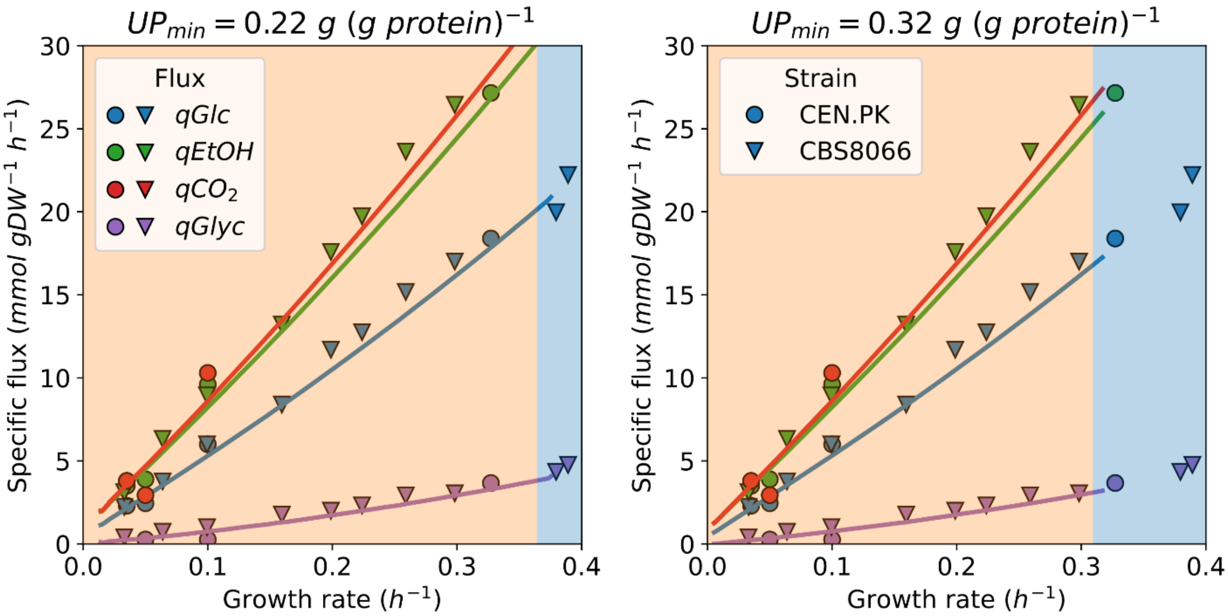


**Supplementary Figure 4. Specific uptake and excretion fluxes as a function of growth rate in glucose-limited chemostat cultures without additional proteomic burden**. In this simulation, the GAM value was set to 40 $mmolgDW^{-1}$ and minimal UP fraction in the proteome to 0.22 $gUP(gprotein)^{-1}$ (left) or 0.32 $gUP(gprotein)^{-1}$ (right). Shading of the panels corresponds to active proteome constraints at different simulation points as represented in Main Text Figure 2. Data for CEN.PK (circles) from (Björkeroth et al., 2020; Jewett et al., 2013; Tai et al., 2007, 2005), and CBS8066 strain (triangles) from (Nissen et al., 1997).

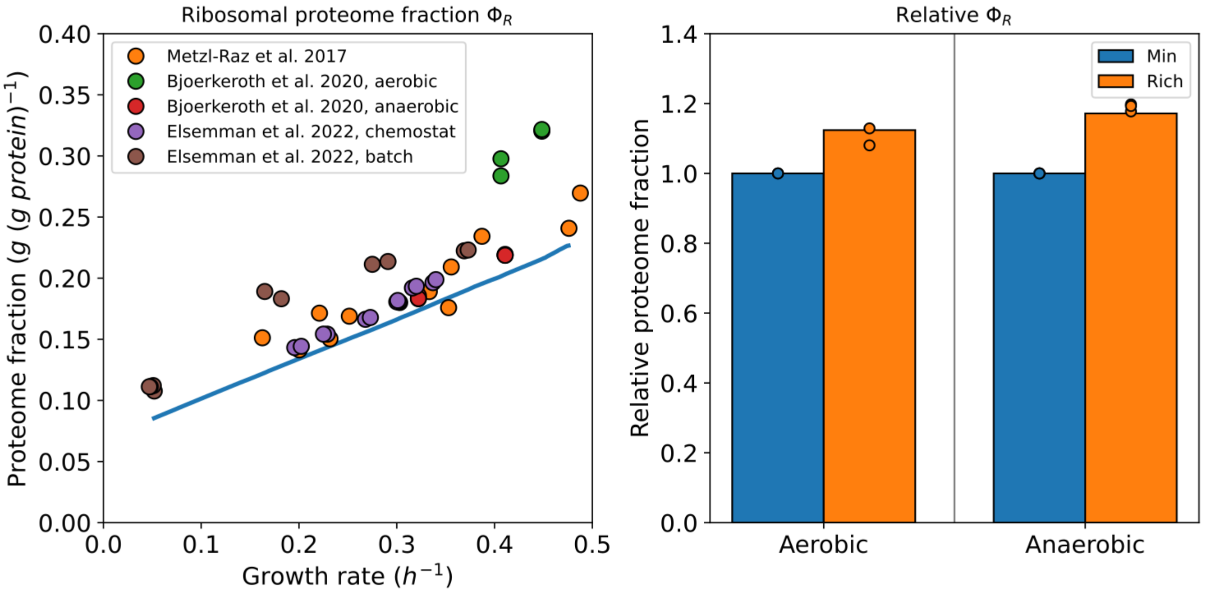


**Supplementary Figure 5. Ribosomal proteome fraction as a function of the growth rate and nutrient upshift. (Left)** ribosomal proteome fraction $\Phi_R$ as a function of the growth rate. Line is model prediction (condition-independent), points are experimental measurements. **(Right)** Relative $\Phi_R$ upon a nutritional upshift from glucose-minimal to rich medium (amino acid supplementation). Data for (left) from (Björkeroth et al., 2020; Elsemman et al., 2022; Metzl-Raz et al., 2017), for (right) from (Björkeroth et al., 2020).

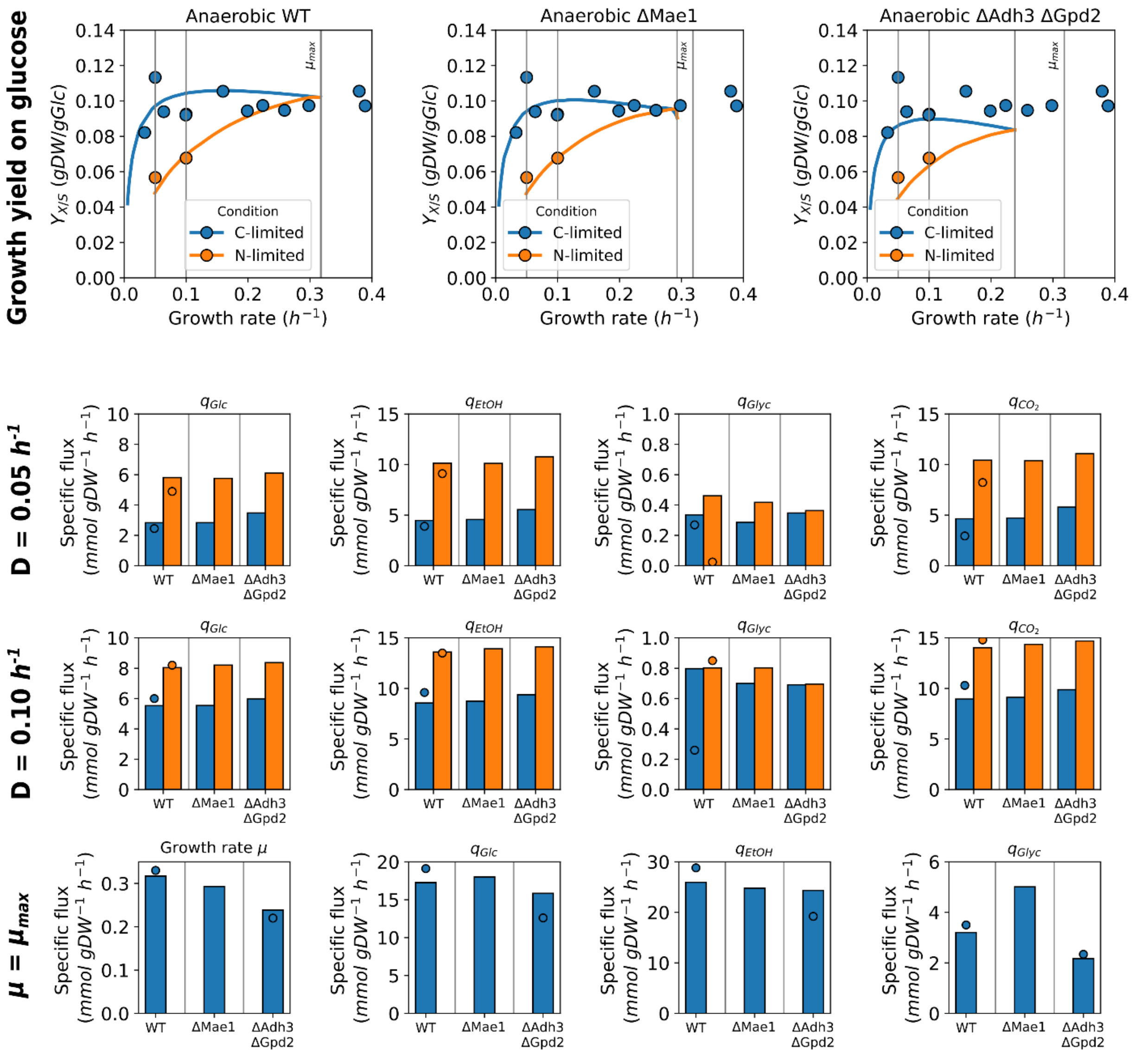


**Supplementary Figure 6. The physiological parameters of deletion strains at different glucose- and ammonium-limited chemostats. Top row**, biomass yield on glucose. In all the panels, the points represent experimental measurements from wild-type strains; lines are model predictions. **Rest of the rows**, specific uptake and excretion fluxes, of anaerobic glucose- and ammonium-limited cultures and batch cultures. Points are experimental measurements, bars are model predictions. Data from (Jewett et al., 2013; Nissen et al., 1997; Tai et al., 2005).

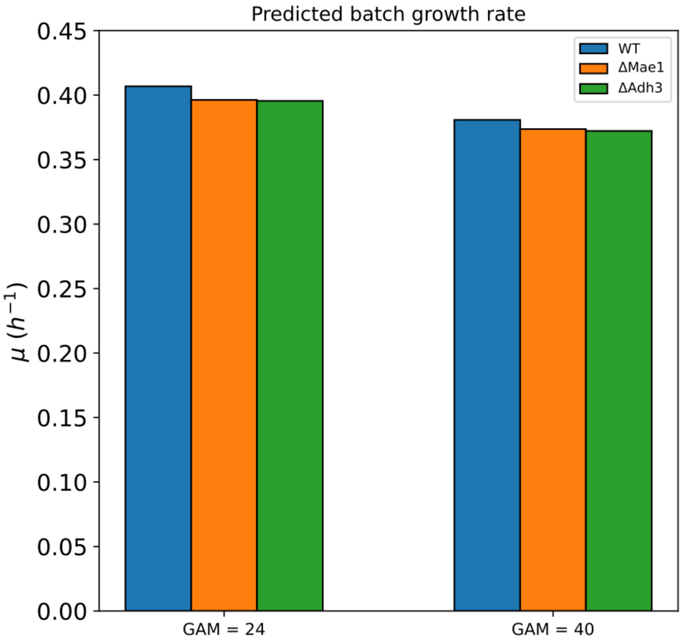


**Supplementary Figure 7. Predicted batch growth rates in amino-acid supplemented media.** The media formulation was as done for Main Text Figure 3 and described in the Methods.

### Supplementary Figures References


Björkeroth, J., Campbell, K., Malina, C., Yu, R., Di Bartolomeo, F., Nielsen, J., 2020. Proteome reallocation from amino acid biosynthesis to ribosomes enables yeast to grow faster in rich media. Proc. Natl. Acad. Sci. 117, 21804–21812. https://doi.org/10.1073/pnas.1921890117

Elsemman, I.E., Rodriguez Prado, A., Grigaitis, P., Garcia Albornoz, M., Harman, V., Holman, S.W., van Heerden, J., Bruggeman, F.J., Bisschops, M.M.M., Sonnenschein, N., Hubbard, S., Beynon, R., Daran-Lapujade, P., Nielsen, J., Teusink, B., 2022. Whole-cell modeling in yeast predicts compartment-specific proteome constraints that drive metabolic strategies. Nat. Commun. 13, 801. https://doi.org/10.1038/s41467-022-28467-6

Jewett, M.C., Workman, C.T., Nookaew, I., Pizarro, F.A., Agosin, E., Hellgren, L.I., Nielsen, J., 2013. Mapping Condition-Dependent Regulation of Lipid Metabolism in *Saccharomyces cerevisiae*. G3 GenesGenomesGenetics 3, 1979–1995. https://doi.org/10.1534/g3.113.006601

Metzl-Raz, E., Kafri, M., Yaakov, G., Soifer, I., Gurvich, Y., Barkai, N., 2017. Principles of cellular resource allocation revealed by condition-dependent proteome profiling. eLife 6, e28034. https://doi.org/10.7554/eLife.28034

Nissen, T.L., Schulze, U., Nielsen, J., Villadsen, J., 1997. Flux Distributions in Anaerobic, Glucose-Limited Continuous Cultures of Saccharomyces Cerevisiae. Microbiology 143, 203–218. https://doi.org/10.1099/00221287-143-1-203

Tai, S.L., Boer, V.M., Daran-Lapujade, P., Walsh, M.C., de Winde, J.H., Daran, J.-M., Pronk, J.T., 2005. Two-dimensional Transcriptome Analysis in Chemostat Cultures. J. Biol. Chem. 280, 437–447. https://doi.org/10.1074/jbc.M410573200

Tai, S.L., Daran-Lapujade, P., Luttik, M.A.H., Walsh, M.C., Diderich, J.A., Krijger, G.C., van Gulik, W.M., Pronk, J.T., Daran, J.-M., 2007. Control of the Glycolytic Flux in Saccharomyces cerevisiae Grown at Low Temperature. J. Biol. Chem. 282, 10243–10251. https://doi.org/10.1074/jbc.M610845200
