## Supplementary Notes for "Elevated energy costs of biomass production in mitochondrial-respiration deficient *Saccharomyces cerevisiae*"

### pcYeast8 model documentation

Pranas Grigaitis *et al.*

November 14, 2022

### Overview

This document summarizes the new version of the proteome-constrained model of *Saccharomyces cerevisiae*, the *pcYeast8*, based on the *Yeast8* metabolic model (version 8.4.2).

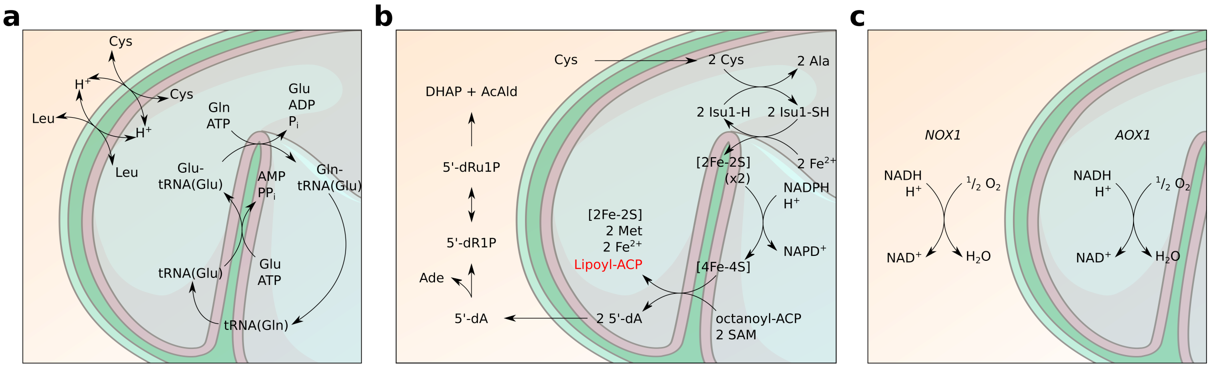

Figure 1: Modifications in the *Yeast8* metabolic model structure. **a.** Amendments to the amino acid turnover processes in mitochondria. **b.** Reconstructed pathway of the lipoic acid biosynthesis. **c.** Localization and reactions of the alternative oxidases *NOX1* and *AOX1*.

### 1 Modifications of the metabolic model

#### 1.1 Amino acid turnover in mitochondria

A central expansion, compared to the conventional genome-scale model, is the introduction of fine-grained descriptions of protein synthesis, folding and degradation. In eukaryotes, ca. 20 proteins are encoded in the mitochondrial genome and translated by the mitochondrial ribosome. Therefore, descriptions of amino acid transport in and out of mitochondria, as well as charging of amino acids onto transport RNAs (tRNAs) should be complete. In the *Yeast8*, however, this was not the case. Therefore we reviewed these processes and included amendments, which we summarized in Figure 1a.

##### Amino acid transport

Transporters for L-cysteine (Cys) and L-leucine (Leu) were missing. We added reactions *r_4705* and *r_4706* for Cys and Leu transport, respectively: AA[c]+$H^+$[c] ↔ AA[m]+$H^+$[m]. Both reactions were set to be catalyzed by amino acid permease *YDR046C*. In addition, all already existing amino acid transporters were set to be reversible.

##### Amino acid-tRNA synthases

For some amino acids, reactions of tRNA charging were missing. These were L-alanine (Ala), Cys, glycine (Gly), and L-serine (Ser). We first added species of both free and charged tRNAs to the model (*s_4263* to *s_4270*) for Ala (free, charged) Cys, Gly, and Ser. Next, we added amino acid-tRNA synthase reactions (*r_4700* to *r_4703*, the same order): AA[m] + tRNA(AA)[m] + ATP[m] → AA-tRNA(AA)[m] + AMP[m] + $PP_i$[m]. We set these reactions to be catalyzed by: *YOR335C* for Ala, *YNL247W* for Cys, *YBR121C* for Gly, and *YHR011W* for Ser.

##### tRNA turnover

In the model, the turnover of a tRNA, specific for L-glutamine (Gln), was poorly designed. We mitigated this problem by:

- Reaction *r_4155* (Glutamyl-tRNA(Gln) amidotransferase subunit B, mitochondrial): add *s_0749* (Glu-tRNA(Glu)) with coefficient −1.0, and delete reagent *s_3754* (L-glutamyl-tRNA(Gln)).
- Create species *s_4271* (tRNA(Gln))
- Create reaction *r_4704*, tRNA(Gln)[m] (*s_4271*) to tRNA(Glu)[m] (*s_1592*)

#### 1.2 Biomass equation

We modified the original biomass equation of the *Yeast8* model (*r_4041*).

##### 1.2.1 Modifications to the original biomass equation

###### Removal of "Growth" reaction

The biomass equation had, among its products, a species called "Biomass" (*s_0450*), which were diluted in a dedicated reaction "Growth" (*r_2111*). Both the species *s_0450* and reaction *r_2111* were deleted from the model. Non-growth-associated maintenance (NGAM) reaction *r_4046* was also removed, as ATP maintenance costs are recomputed at every step of the simulation.

###### Removal of metabolite pools

The model will explicitly account for proteome turnover, thus the biomass equation itself should not include dilution of proteins by growth. Moreover, some biomass components, like cofactors or ions, are of little influence to the outcome of model simulations. Therefore we removed the following reagents from the reaction *r_4041* for the sake of analyzing the model results more easily:

- Protein pool (*s_3717*)
- Cofactor pool (*s_4205*)
- Ion pool (*s_4206*)

###### Specifying non-mitochondrial lipid composition

*Yeast8* introduced the SLIMER formalism for defining lipid composition in the biomass. However, instead we used lipids measurements of (Ejsing et al., 2009) to modify the lipid pseudoreaction (*r_2108*). First, we deleted the substrates of this reaction, and then added lipid species quantified by Ejsing and colleagues as substrates with stoichiometric coefficients, representing $mmol\ (g\ lipid)^{-1}$ of each lipid (a flux of 1 through *r_2108* would mean 1 gram of lipids produced). As we defined a separate mitochondria-specific biomass equation, this equation does not include mitochondria-specific lipids.

##### 1.2.2 Growth rate-dependent biomass composition

The biomass composition of the CEN.PK113-7D strain changes in a growth rate-dependent manner, as reported by (Lange and Heijnen, 2001; Canelas et al., 2011). Therefore, at every iteration of the binary search, we recompute the coefficients for biomass components based on the growth rate. A summary of the growth rate-relations of different biomass components can be found in Table 1.

#### 1.3 Lipoic acid synthesis and 5'-deoxyadenosine salvage pathway

Lipoic acid is an essential cofactor for mitochondrial metabolism: it plays a role in catalytic processes of pyruvate dehydrogenase complex, $\alpha$-ketoglutarate dehydrogenase and other enzymes. The precursor of lipoic acid synthesis is octanoic acid (C8:0), bound to the ACP (acyl carrier protein). In the final step of the synthesis, a disulphide bridge is formed at $C_{7-8}$ position (reviewed in more depth in (Solmonson and DeBerardinis, 2018)). However, the *Yeast8* model does include a generic description of lipoic acid synthesis (*r_4324*), containing two dead-end metabolites ("sulfur donor", *s_4004* and "5'-deoxyadenosine", *s_4005*). Presence of dead-end metabolites means that any flux through reaction *r_4324* will result in violation of the steady-state condition. We therefore reconstructed two pathways in order for model to produce lipoic acid. First we introduced biosynthesis of [Fe-S] clusters, which act as sulfur donors in lipoic acid synthesis. Moreover, we added the 5'-deoxyadenosine salvage pathway to the model, based on literature data.

###### [Fe-S] cluster biosynthesis

Instead of using a generic "sulfur donor", we implemented the reactions needed for generation of [4Fe-4S] clusters, needed for lipoic acid biosynthesis. We largely followed (Braymer and Lill, 2017; Lill et al., 2020) for designing the process. First, two new genes ("YLL027W", "YPR067W") were added to the model, as well as two [Fe-S] cluster species: "[2Fe-2S] cluster" and "[4Fe-4S] cluster".

Table 1: Growth rate-dependent biomass composition. Data collected from (Lange and Heijnen, 2001; Canelas et al., 2011; Tada and Tada, 1962)

| Component | Reaction | Species | Relation |
|---|---|---|---|
| **Lipids** | *r_4041* | *s_1096* | $m_{Lipids} = -0.045576 \times \mu + 0.074549$ |
| **Carbohydrates** | | | |
| Trehalose | *r_4048* | *s_1520* | $12.61369 \times exp(-13.10818\mu)/100.0) \times 1e3/342.3$ |
| Glycogen | *r_4048* | *s_0773* | $20.44753 \times exp(-9.593868\mu)/100.0) \times 1e3/162.0$ |
| $\beta-$glucans | | | $m_{glucans} = -0.137727\mu + 0.188208$ |
| $(1 \rightarrow 3)$ | *r_4048* | *s_0001* | $m_{glucans}/2.0 \times 1e3/162.0$ |
| $(1 \rightarrow 6)$ | *r_4048* | *s_0004* | $m_{glucans}/2.0 \times 1e3/162.0$ |
| Mannans | *r_4048* | *s_1107* | $0.09975 \times 1e3/162.0$ |
| **RNA** | | | $m_{RNA} = 0.15249 \times \mu + 0.049841$ |
| AMP | *r_4049* | *s_0423* | $m_{RNA} \times 1e3 \times 0.21626/347.22$ |
| GMP | *r_4049* | *s_0782* | $m_{RNA} \times 1e3 \times 0.32011/363.22$ |
| CMP | *r_4049* | *s_0526* | $m_{RNA} \times 1e3 \times 0.26167/323.1965$ |
| UMP | *r_4049* | *s_1545* | $m_{RNA} \times 1e3 \times 0.20190/324.1813$ |
| **Protein** | | | $m_{Prot} = 0.42436 \times \mu + 0.35364$ |
| **ATP** | | | |
| GAM part | *r_4041* | | $24.0\ mmol\ gDW^{-1} \times \mu$ |
| NGAM part | *r_4041* | | $0.168\ mmol\ gDW^{-1}\ h^{-1}$ |

###### 5'-deoxyadenosine salvage pathway

A product of lipoic acid synthesis, 5'-deoxyadenosine, has to be metabolized into end-products acetaldehyde and dihydroxyacetone phosphate (DHAP). Following (Rapp and Forchhammer, 2021), we added species "5'-Deoxyadenosine' (cytosolic species), "5'-deoxy-D-ribose 1-phosphate", and "5'-deoxy-D-ribulose 1-phosphate, as well as reactions needed for this process (all genes needed were present in the model already).

###### Modifications to the lipoic acid synthesis reaction

We accordingly modified the reaction *r_4324*, the last step of lipoic acid biosynthesis pathway. First, we removed reagent "sulphur donor" (*s_4004*). Instead of it, we added reagents from the [Fe-S] cluster biosynthesis pathway. The final reconstructed pathway is shown in Figure 1b.

#### 1.4 Mitochondria-specific maintenance reaction

In order to more precisely reflect on the costs, specific to the mitochondria, we added an additional reaction to the model, $v_{MitoMaintenance}$. Substrates of this reaction are mitochondria-specific lipids, as well as mitochondrial ATP (Table 2). Unlike the biomass equation, the stoichiometric coefficients in this equation are scaled to gram of mitochondrial protein, rather than gram of dry cell weight. As the abundances of individual mitochondrial proteins are optimization variables, so is the flux through this reaction. Therefore, we set the flux through this reaction using an additional coupling constraint, dependent on the mitochondrial protein content (see Section 3.1.2).

#### 1.5 Alternative NADH oxidases

In the light of experiments of (Vemuri et al., 2007), we also wanted to investigate the effect to model predictions of overflow metabolism by two heterologous water-forming NADH oxidases, *aox* from *Ajellomyces capsulatus* (UniProt ID: Q9Y711) and *nox* from *Streptococcus pneumoniae* (UniProt ID: O84925). Experimentally, *nox* was a cytosolic protein, while *aox* was targeted to mitochondria (Figure 1c). Thus we attributed gene IDs *AOX1* (for *aox*) and *NOX1* (for *nox*) and added these reactions, with respective gene-protein-reaction (GPR) associations, to the model (*r_4711* for *AOX1* and *r_4712* for *NOX1*).

Table 2: Components of mitochondrial maintenance reaction as a function of mitochondrial proteome mass $m_{MitoProtein}$. Data collected from (Blagović et al., 2005; Zinser et al., 1991; Zinser and Daum, 1995)

| Component | Species | Stoich. coefficient |
|---|---|---|
| **Lipids** | | |
| cardiolipin (1-16:0, 2-18:1, 3-18:0, 4-16:1) | *s_3260* | -0.000408333 |
| cardiolipin (1-16:1, 2-16:1, 3-16:0, 4-16:1) | *s_3240* | -0.001434249 |
| cardiolipin (1-16:1, 2-16:1, 3-16:1, 4-16:1) | *s_3241* | -0.001434249 |
| cardiolipin (1-18:0, 2-16:1, 3-18:0, 4-16:1) | *s_3248* | -0.001434249 |
| cardiolipin (1-18:1, 2-18:1, 3-16:0, 4-18:1) | *s_3312* | -0.001434249 |
| ergosterol | *s_0667* | -0.025211247 |
| phosphatidate (1-16:0, 2-16:1) | *s_3092* | -0.002485448 |
| phosphatidate (1-16:0, 2-18:1) | *s_3103* | -0.000130860 |
| phosphatidate (1-18:0, 2-16:1) | *s_3099* | -0.000157566 |
| phosphatidyl-L-serine (1-16:0, 2-16:1) | *s_3128* | -0.003285675 |
| phosphatidyl-L-serine (1-16:0, 2-18:1) | *s_3137* | -0.000173850 |
| phosphatidyl-L-serine (1-18:0, 2-16:1) | *s_3133* | -0.000236350 |
| phosphatidylcholine (1-16:0, 2-16:1) | *s_3296* | -0.043928864 |
| phosphatidylcholine (1-16:0, 2-18:1) | *s_3304* | -0.002324110 |
| phosphatidylcholine (1-18:0, 2-16:1) | *s_3300* | -0.002798418 |
| phosphatidylethanolamine (1-16:0, 2-16:1) | *s_3130* | -0.031459506 |
| phosphatidylethanolamine (1-16:0, 2-18:1) | *s_3138* | -0.001660669 |
| phosphatidylethanolamine (1-18:0, 2-16:1) | *s_3134* | -0.001999581 |
| Lipoyl-[acp] | s_3946 | -0.000180995 |
| ACP1 | s_1845 | 0.000180995 |
| **ATP** (hydrolysis) | | -6.0 |

#### 1.6 Minor fixes

##### Modified species IDs

In the *Yeast8* model, species IDs are constructed from so-called "Replacement IDs" and compartment codes. However, the SBML formulation does not allow "[]" to be in any of the objects' IDs and thus they are replaced by "___91___" and "___93___", for "[" and "]", respectively. This notation complicates readability of the SBML document, as well as model handling. Thus the species IDs were modified so that only the "Replacement IDs" remain (format: "*s_XXXX*", *XXXX* is the 4-digit code for the species).

##### Modified gene IDs

Analogously, gene IDs were modified to represent the ORF names without a dash (e.g. *YDR322C-A* is presented in the model as *YDR322C___45___A*). Substitution was made to get a consistent gene name nomenclature without the dashes (in this example, the final outcome is *YDR322CA*).

##### Compartment dimensions

Compartments were highlighted by CBMPy as "zero-dimension". Therefore, dimensions were added:

- 3 dimensions (default)
- 2 dimensions for membrane compartments and *ce* (cell envelope)

##### Stoichiometry of mitochondrial ATP synthase

In the *Yeast8* model, proton stoichiometry in ATP synthesis is defined incorrectly ($3H^+[c] \rightarrow 2H^+[m]$), while experimentally determined stoichiometry is $4H^+[c] \rightarrow 3H^+[m]$. This was changed in reaction *r_0226* accordingly.

###### Reversibility of glycerol:$H^+$ symporter

In the *Yeast8* model, the mode of glycerol transport through plasma membrane depends on the direction: the uptake of glycerol to the cell was modeled as glycerol:$H^+$ symport (*r*_1171) (Ferreira et al., 2005), and its export was set as uniport "via channel" (*r*_1172). Both of these reactions were irreversible, suggesting that only one mode is active. We set the glycerol:$H^+$ symport reaction (*r*_1171) to be reversible instead.

###### Stoichiometry of monosaccharide import

In *S. cerevisiae*, most of the monosaccharides are known to be imported via facilitated diffusion (a rule of thumb for oligosaccharides is sugar:$H^+$ symport). In the *Yeast8* model, transport of fructose (*r*_1134) and galactose (*r*_1135) were incorrectly set as sugar:$H^+$ symport. We removed $H^+$ species from these reactions.

##### 1.7 Splitting of reactions

###### Splitting `OR` gene-protein-reaction associations

If a gene-protein-reaction (GPR) association did not have an `OR` relationship, then we just used the current GPR association in order to update the reaction name (added the required protein names as a suffix to the reaction name, reaction name *R1* becomes *R1___GPR*). If the reaction GPR had an `OR` relationship, the reaction was deep-copied and new reactions with individual GPR strings were created. E.g.: GPR of a reaction *R1* `(G1 OR (G2 and G3) OR G4)` was split into reactions *R1___G1*, *R1___G2*, *R1___G3*, and *R1___G4*.

###### Splitting reversible reactions

In the *pc*-models, reactions which have proteins associated to them must be unidirectional (otherwise, a reverse reaction would be either impossible and/or would produce metabolic enzymes). Here we split such reactions into two forward and reverse reactions. E.g.: a reversible reaction *R1* was split into forward and reverse reactions *R1_fwd* and *R1_rev*, respectively.

#### 2 Description of proteome turnover and reaction coupling

We created different model species for each protein entity of the UniProt reference proteome of *S. cerevisiae* (*UP000002311*) (UniProt Consortium, 2020) in the model, for a sample species with the UniProt accession ID *P00000*:

- Unfolded protein species (*P00000_uf_c*)
- Folded protein species (*P00000_c*)
- "Used" protein species (*P00000_used_c*)

Although the target compartment in the example is cytoplasm (species suffix "_*c*"), for proteins, translated in mitochondria, or transported to compartments other than cytoplasm, we created corresponding model species as well. The turnover of proteins in the model (conversions from one species to another) can be described through several classes of reactions, which we will describe in the next sections.

For implementing enzyme coupling constraints, we use protein "complex" species. These species represent the enzymatic complexes (could be a single protein as well) at their stoichiometric ratios (if known). A complex formation reaction of $cplx_i$, $v_{cplx\ formation,\ i}$ takes a general form, where complex is formed from individual proteins:

$$v_{cplx\ formation,\ i} : \sum_{j \in cplx_i} n_j\ j\_c \rightarrow cplx_i \tag{1}$$

The flux through $v_{cplx\ formation,\ i}$ is used for coupling with the reactions which need $cplx_i$ for catalysis. To maintain the steady-state assumption, two options of consuming the complex are provided:

degradation of the complex into individual proteins (Eq. 2), and dilution by growth (Eq. 3). These two processes, respectively, are represented as follows:

$$v_{cplx\ degradation,\ i} : cplx_i \rightarrow \sum_{j \in cplx_i} n_j\ j\_used\_c \tag{2}$$

$$v_{cplx\ dilution,\ i} : cplx_i \rightarrow \emptyset \tag{3}$$

The ratio between the flux through complex degradation and dilution-by-growth reactions is growth rate $\mu$-dependent, and is defined in the following section.

#### 2.1 General form of coupling constraints

The use of macromolecular complexes (alike metabolic enzymes and protein turnover machinery [e.g., ribosomes]) is coupled with the fluxes through reactions they catalyze. We describe this relationship between enzyme demand and metabolic flux as follows:

$$v = [e] \times k_{cat} \times f(\mathbf{x}, T, \dots) \tag{4}$$

Where $f$ is a saturation function (ranging [0; 1]), dependent on different parameters, such as metabolite concentrations, temperature, and others. In the *pc*-models, we assume $f = 1$, i.e. enzymes working at their maximal speed. This way, we predict the *minimal* enzyme demand, needed to sustain the metabolic flux. For each enzyme complex, we then couple the flux through complex formation reaction to the sum of fluxes, which that complex catalyzes, scaled with the $k_{cat}$ values:

$$\frac{v_{cplx\ formation,\ i}}{k_{deg} + \mu} = \sum_j \frac{v_j}{k_{cat,\ i,\ j}} \tag{5}$$

Then, we set another two constraints for each complex, describing the degradation of proteins (degraded with rate $k_{deg}$) and dilution by growth (with rate $\mu$):

$$v_{cplx\ degradation,\ i} = v_{cplx\ formation,\ i} \times \frac{k_{deg}}{k_{deg} + \mu} \tag{6}$$

$$v_{cplx\ dilution,\ i} = v_{cplx\ formation,\ i} \times \frac{\mu}{k_{deg} + \mu} \tag{7}$$

#### 2.2 Protein translation

In the model, we describe protein translation as use of loaded amino acid-tRNAs ($tRNA^{AA}(AA)$) and retrieval of free tRNAs ($tRNA^{AA}$). First, free amino acids (AAs) are loaded onto respective tRNAs (Eq. 8), with a cost of 2 equivalents of ATP per amino acid. Then, loaded amino acid-tRNAs are used for protein translation according to the amino acid composition of the example protein *P00000* ($v_{syn,\ P00000}$) (Eq. 9), with $l_{P00000}$ being the peptide chain length. One cycle of peptide chain elongation costs 2 GTP, making sum energetic cost of protein synthesis to be 4 NTPs per amino acid.

$$AA + tRNA^{AA} + ATP \rightarrow tRNA^{AA}(AA) + AMP + PP_i \tag{8}$$

$$v_{syn,\ P00000} : n_1\ tRNA^{AA_1}(AA_1) + \dots + n_{20}\ tRNA^{AA_{20}}(AA_{20}) + 2l_{P00000}\ GTP + 2l_{P00000}\ H_2O \rightarrow$$
$$n_1\ tRNA^{AA_1} + \dots + n_{20}\ tRNA^{AA_{20}} + 2l_{P00000}\ GDP + 2l_{P00000}\ P_i + 2l_{P00000}\ H^+ + P00000\_uf\_c \tag{9}$$

###### Ribosome capacity coupling

We described the coupling between protein synthesis fluxes and ribosome formation as a function of peptide length. The ribosomal peptide elongation rate $k_{cat,\ ribo} = 10.5\ aa\ s^{-1} = 3.78 \times 10^4\ aa\ h^{-1}$ (Metzl-Raz et al., 2017) is used to scale the ribosome demand to the protein synthesis fluxes. Also, (Metzl-Raz et al., 2017) have showed that the fraction of ribosomal proteins in the total proteome increases linearly with the growth rate $\mu$, with a offset value of $\phi_R^0 \approx 0.08\ g\ (g\ protein)^{-1}$. As discussed in the Supplementary Note 6 of the pcYeast7.6 model (Elsemman et al., 2021), in order to capture the offset in the model, we need to decompose this value into respective values for (a) truly inactive ribosomes $\phi_R^{0'}$ and (b) the ribosomes, required to synthesize the inactive ribosomes. Previously, we reported the fraction $\phi_R^{0'} \approx 0.053$ (Elsemman et al., 2021). With the molecular weight of the protein component of the ribosome $MW_{P,\ ribo} = 1388.488\ g\ mmol^{-1}$, we compute the increment in the flux through ribosome complex formation reaction. Thus the general formulation of coupling constraint in Eq. 5 becomes:

$$\frac{k_{cat,\ ribo}[aa\ h^{-1}]}{k_{deg} + \mu} \times v_{cplx\ formation,\ ribosome} = \sum_{i \in cytoRibo} l_i[aa] \times v_{syn,\ i} + \frac{\phi_R^{0'} \times m_{prot}}{MW_{P,\ ribo}} \tag{10}$$

##### Translation factor demand

A number of translation initiation, elongation, and termination factors are needed for protein synthesis in ribosomes (Table 3). We thus considered different aspects of the translation for these groups of translation factors.

We first considered the translation initiation factors (eukaryotic initiation factors, eIFs). Their demand can be linked with the time, needed for ribosome to locate the initiation codon and fully assemble. For this, we first collected information on the length of the 5'-untranslated region (5'-UTR) from Saccharomyces genome database (SGD, (Cherry et al., 2012)). For mRNAs without a determined length of the 5'-UTR, we assumed the median value of 60 nucleotides. We assumed that ribosome scans the 5'-UTR region of the mRNA at the speed of $k_{cat,\ eIF} = 10\ nt\ s^{-1} = 3.6 \times 10^4\ nt\ h^{-1}$. Following that, we set the following coupling constraints for each of the individual complexes of translation initiation factors (Table 3):

$$\frac{k_{cat,\ eIF}[nt\ h^{-1}]}{k_{deg} + \mu} \times v_{cplx\ formation,\ eIF_i} = \sum_{j \in cytoRibo} l_{5'UTR,\ j}[nt] \times v_{syn,\ j} \tag{11}$$

Next, we coupled the demand for translation elongation factors (eEFs) in a similar manner. It should be noted that clear estimates of the turnover values of these factors are not available. Thus we reason that the $k_{cat,\ EF}$ value should be $k_{cat,\ eEF} \geq 10.5\ aa\ s^{-1}$. We can assume this because otherwise, the observed maximal speed of peptide elongation $k_{cat,\ ribo}$ would be considerably lower, as elongation is the longest phase in the translation process. Using this information, we set the coupling constraints for all individual complexes of translation elongation factors:

$$\frac{k_{cat,\ eEF}[aa\ h^{-1}]}{k_{deg} + \mu} \times v_{cplx\ formation,\ eEF_i} = \sum_{j \in cytoRibo} l_j[aa] \times v_{syn,\ j} \tag{12}$$

Eukaryotic release factor (eRF) is the protein complex (eRF1 and eRF3), facilitating the dissociation of ribosomes (termination of translation) and release of fully translated peptide. The termination rate is measured to be $k_{cat,\ eRF} = 0.15\ s^{-1} = 540\ h^{-1}$ (Shoemaker and Green, 2011).

$$\frac{k_{cat,\ eRF}[h^{-1}]}{k_{deg} + \mu} \times v_{cplx\ formation,\ eRF} = \sum_{i \in cytoRibo} v_{syn,\ i} \tag{13}$$

##### Note on mitochondrial translation

Mitochondria also have their own ribosomes, which translate a handful of protein species (including one subunit of the mitochondrial ribosomes). We set coupling constraints, similar to the cytosolic ribosomes, to describe the demand of mitochondrial ribosomes:

$$\frac{k_{cat,\ mito\ ribo}[aa\ h^{-1}]}{k_{deg} + \mu} \times v_{cplx\ formation,\ mito\ ribosome} = \sum_{i \in mitoRibo} l_i[aa] \times v_{syn,\ i} \tag{14}$$

We chose to estimate the $k_{cat,\ mito\ ribo}$ value, rather than apply a "offset" value, as we did for cytosolic ribosomes. We thus fitted the peptide elongation rate of mitochondrial ribosomes, using the proteomics

Table 3: Translation factors (and groups of), defined in the pcYeast8 model.

| Factor type | Cytosolic factors | Mitochondrial factors |
|---|---|---|
| Initiation factors | eIF1, eIF1A<br>eIF2, eIF2A, eIF2B<br>eIF3<br>eIF4A, eIF4B, eIF4E, eIF4F<br>eIF5, eIF5A, eIF5B<br>eIF6 | mIF |
| Elongation factors | eEF1A, eEF1B, eEF1G<br>eEF2<br>eEF3A, eEF3B | mEF |
| Release factors | eRF | mRF |

data from our previous study (Elsemman et al., 2021). The value of $k_{cat,\ mito\ ribo} = 8\ aa\ s^{-1} = 2.8 \times 10^4\ aa\ h^{-1}$ has shown the best agreement to the experimental measurements. We also set up constraints to describe the use of mitochondrial translation factors, with similar expressions (Eqs. 11, 12, and 13) the same $k_{cat}$ values as for their cytosolic counterparts.

#### 2.3 Protein folding and transport

The unfolded protein species (synthesis described by Eq. 9) then have to be folded into "folded" (active) protein species. Optionally, these proteins first have to be transported to their target compartments. The folding reactions $v_{folding,\ P00000}$, catalyzed by chaperones (see coupling constraints, explained later), take the following form:

$$v_{folding,\ P00000} : P00000\_uf\_c \rightarrow P00000\_c \tag{15}$$

The transport reactions (transport demands explicitly modelled for mitochondria, through use of TIM/TOM translocase system) for the unfolded protein species **to** the target compartment (e.g. Golgi) and for the used protein species **out** of the compartment (except for mitochondria, see "Protein degradation") are as follows:

$$v_{import,\ P00000} : P00000\_uf\_c \rightarrow P00000\_uf\_g \tag{16}$$

$$v_{export,\ P00000} : P00000\_used\_g \rightarrow P00000\_used\_c \tag{17}$$

##### Chaperone coupling

We specified some chaperones in the model (Table 4) for protein folding. Some of these chaperones are specific for some compartments (such as BiP or mitochondrial Hsp class chaperons), others were set to work in other compartments. Given that several chaperones could fold the same clients, we also attributed arbitrary coefficients $c_i$ ("workload sharing") to different chaperones, based on their abundance in proteome. The coefficients of mitochondrial chaperones, unlike their counterparts in other compartments (Table 4), add up to 2 due to the demand of chaperone action during translocation to mitochondria and degradation (see next subsections). In the end, similarly to coupling constraints for ribosomes or translation factors, we described the coupling between folding reactions and chaperone use as follows:

$$\frac{k_{cat,\ chap_i}[h^{-1}]}{k_{deg} + \mu} \times v_{cplx\ formation,\ chap_i} = c_i \sum_{j \in chap_i} v_{folding,\ j} \tag{18}$$

##### TIM/TOM translocase coupling

We modeled protein transport to mitochondria explicitly due to the fact that mitochondria are very metabolically active organelles and also they impose proteome limits on cellular metabolism.
Translocation of proteins to mitochondria begins by pulling the unfolded peptide through the channels, formed by the outer- and inner membrane translocases (denoted as $t_i$ in equations). Mitochondrial (mt) mtHsp60 and mtHsp70 play an important role in these processes. Importantly, mtHsp70 pulls the

Table 4: Chaperones, defined in the pcYeast8 model.

| Chaperone | Coefficient | Target compartment(s) | $k_{cat}[h^{-1}]$ | Source |
|---|---|---|---|---|
| SSA<br>SSB<br>CCT<br>PFD<br>Hsp90<br>Hsp104 | 0.25<br>0.25<br>0.125<br>0.125<br>0.125<br>0.125 | Cytosol<br>Nucleus<br>Vacuole<br>Peroxisome | 57.0 | Lopez-Buesa et al. (1998)<br>(for SSA/SSB) |
| BiP | 1.0 | Golgi<br>ER | 78.0 | Rosam et al. (2018) |
| mtHsp60<br>mtHsp70<br>mtHsp78 | 0.5<br>0.5<br>1.0 | Mitochondria | 57.0 | Same as for SSA/SSB |

unfolded peptide through the translocases, using ATP hydrolysis energy (Okamoto et al., 2002), and both mtHsp60 and mtHsp70 can fold the translocated peptide. Three kinds of translocases are defined in the model: TOM, the outer membrane translocase, TIM22, and TIM23 (both inner membrane translocases). We estimated the turnover value of $k_{cat,\ t} = 3\ min^{-1} = 180\ h^{-1}$, and set the following coupling constraint for each translocase complex:

$$\frac{k_{cat,\ t}[h^{-1}]}{k_{deg} + \mu} \times v_{cplx\ formation,\ t_i} = \sum_{j \in mito} v_{import,\ j} \tag{19}$$

#### 2.4 Protein degradation

Used protein species are degraded to individual amino acids either in cytosol (cytosolic proteins and proteins from compartments other than mitochondria) or in mitochondria. In the first case, degradation is catalyzed by the proteasome complex, in the latter - by mitochondrial protease *PIM1*. In both cases, proteins are hydrolyzed to amino acids, consuming an estimated 1.35 ATP per amino acid released (Hong et al., 2012).

$$\begin{aligned} v_{degradation,\ P00000} : P00000\_used\_c + 1.35n\ ATP + 2.35n\ H_2O \rightarrow \\ n_1\ AA_1 + \cdots + n_{20}\ AA_{20} + 1.35n\ ADP + 1.35n\ P_i + 1.35n\ H^+ \end{aligned} \tag{20}$$

##### Proteasome and *PIM1* coupling

As mentioned in the section above, all but mitochondria-targeted proteins are exported as their used forms from their target compartments and degraded in the cytosol (Eq. 17). In the model, we couple cytosolic protein degradation with the use of proteasomes. A proteasome is a large protein complex which unfolds and degrades proteins into short peptides and/or free amino acids. Similarly, in mitochondria, both unfolding and degradation happen, yet catalyzed by two separate proteins: mtHsp78 is responsible for protein unfolding, and PIM1 is the homolog of prokaryotic *Lon* protease. We considered the turnover values of both proteasome $k_{cat,\ proteasome} = 2.3\ min^{-1} = 138\ h^{-1}$ (Peth et al., 2013) and PIM1 $k_{cat,\ PIM1} = 0.26\ s^{-1} = 936\ h^{-1}$ (Patterson-Ward et al., 2007) from literature, and set the coupling constraints in both cases:

$$\frac{k_{cat,\ proteasome}[h^{-1}]}{k_{deg} + \mu} \times v_{cplx\ formation,\ proteasome} = \sum_{j \in proteasome} v_{degradation,\ j} \tag{21}$$

and

$$\frac{k_{cat,\ mtHsp78}[h^{-1}]}{k_{deg} + \mu} \times v_{cplx\ formation,\ mtHsp78} = \sum_{j \in PIM1} v_{degradation,\ j} \tag{22}$$

$$\frac{k_{cat,\ PIM1}[h^{-1}]}{k_{deg} + \mu} \times v_{cplx\ formation,\ PIM1} = \sum_{j \in PIM1} v_{degradation,\ j} \tag{23}$$

### 3 Compartment-specific proteome constraints

In the model, we use three sets of different compartment-specific proteome constraints (also called "capacity constraints"), by the parameter of the proteins which is used to formulate the constraint: protein mass (molecular weight), protein crossection surface area, and protein volume. In the following, we describe these constraints in more detail.

#### 3.1 Proteome mass constraints

##### 3.1.1 Total proteome mass constraint

The proteome mass constraint is an equality constraint, i.e. its expression must always satisfy the right-hand-side (RHS). This constraint therefore sets the protein density in the dry biomass to be consistent with experimental measurements. For the *S. cerevisiae* strain we collected the majority of the data, CEN.PK 113-7D, (Canelas et al., 2011) has determined that the bulk protein mass fraction the dry weight $f_{p,biomass}(\mu)$ increases with increasing growth rate $\mu$ (Table 1). Following these measurements, we set the total proteome mass to equal:

$$\sum_i \frac{v_{syn,i}}{k_{deg} + \mu} \times MW_i \ [\frac{g\ protein}{mol}] \times 10^{-3} \ [\frac{mol}{mmol}] = f_{p,biomass}(\mu) \ [(g\ protein)\ gDW^{-1}] \tag{24}$$

##### 3.1.2 Mitochondrial maintenance coupling

We described the mitochondria-specific maintenance reaction (Section 1.4), which we used to describe the costs of maintaining mitochondria, not explicitly included in the model. The energetic costs in the maintenance reaction also account for the ATP hydrolysis, used in translocation of proteins into mitochondria (Section 2.3). All the components in the mitochondria-specific biomass equation were specified per gram mitochondrial protein, thus we coupled the flux through the maintenance reaction to the mass of mitochondrial proteins:

$$\sum_{i \in mito} \frac{v_{syn,i}}{k_{deg} + \mu} \times MW_i \ [\frac{g\ protein}{mol}] \times 10^{-3} \ [\frac{mol}{mmol}] = v_{Mitochondrial\ maintenance} \tag{25}$$

##### 3.1.3 Unspecified protein constraint

In the model, not all proteins, present in the proteome of *S. cerevisiae* are defined. These proteins can be without enzymatic annotation (e.g. structural proteins), not directly metabolic (e.g. signaling proteins) or simply be not annotated. However, our previously published proteomics data (Elsemman et al., 2021) suggests that more than 2-2.5 thousand protein species can be robustly quantified using label-free mass spectrometry-based proteomics, which is a substantially higher number than the number of protein species, predicted to be expressed by the model. In order to account for the protein expression, which are not explicitly defined in the model, we created an artificial protein species, *PDUMMY*, which we call the "unspecified protein" (UP). Based on the proteomics measurements in glucose-limited chemostats, proteins that are not represented in the model, occupy ca. 25% of the total protein mass. Thus we set the minimal level of expression of the UP at every growth rate $\mu$, which corresponds to the proteome mass fraction $\phi_{UP}$. Note that we specified some maintenance requirements for the mitochondria. Following suit, the unspecified protein demand thus was also split into two constraints, one for the cytosolic ("general") UP expression demand:

$$\frac{v_{folding,\ cyto,\ UP}}{k_{deg} + \mu} \times MW_{UP} \ [\frac{g\ UP}{mol}] \times 10^{-3} \ [\frac{mol}{mmol}] \geq \phi_{UP} \ [(g\ UP)\ (g\ protein)^{-1}] \times f_{p,biomass}(\mu) \tag{26}$$

Alternatively, can we formulate the (Eq. 26) so that we do not have to explicitly input the value of $f_{p,biomass}(\mu)$ into the constraint expression:

$$\sum_{i \in proteins} \frac{v_{syn,\ i}}{k_{deg} + \mu} \times MW_i \ [\frac{g\ protein}{mol}] - \frac{1}{1 - \phi_{UP}} \times MW_{UP} \ [\frac{g\ UP}{mol}] \times v_{folding,\ cyto,\ UP} \leq 0 \tag{27}$$

Then, the mitochondrial UP experession demand (determined to be 28% of total mitochondrial proteome) is set as follows:

$$\sum_{i \in mito\ proteins} \frac{v_{folding,\ mito,\ i}}{k_{deg} + \mu} \times MW_i\ [\frac{g\ protein}{mol}] - \frac{1}{1 - \phi_{UP,\ mito}} \times MW_{UP}\ [\frac{g\ UP}{mol}] \times v_{folding,\ mito,\ UP} \leq 0 \tag{28}$$

##### 3.1.4 Gratuitous protein expression constraint

As one of the tests, we aimed to capture the effects of gratuitous protein expression on the growth rate. Following the quantitative measurements of (Kafri et al., 2016), for this we have specified a gratuitous protein *mCherry* in the model. Similarly to the UP expression constraint (Eq. 26), we can define the proteome mass fraction of the mCherry $\phi_{mCherry}$:

$$\frac{v_{syn,mCherry}}{k_{deg} + \mu} \times MW_{mCherry}\ [\frac{g\ mCherry}{mol}] \times 10^{-3}\ [\frac{mol}{mmol}] = \phi_{mCherry} \times f_{p,biomass}(\mu) \tag{29}$$

#### 3.2 Protein area and volume constraints

**Constants and bionumbers for conversion**

These constraints are expressed by area/volume per cell, therefore, some conversion units are needed for establishing relations between parameters of proteins and cells.
First, we compute the volume of a protein molecule, based on its molecular weight. Following (Erickson, 2009), we assume the protein to be globular (to form a sphere when folded). Then, we compute the minimal radius of the protein molecule with that molecular weight ($r_{min}$), using the following empirical relation:

$$r_{min}\ [nm] = 0.066 \times MW^{\frac{1}{3}} \tag{30}$$

Further, assuming the sphere-shaped protein molecule, we can compute its crossection area (area of a disc with the radius $r_{min}$) and volume (also using the radius $r_{min}$):

$$A_{peptide}\ [nm^2] = \pi \times r_{min}^2 \tag{31}$$

$$V_{peptide}\ [nm^3] = \frac{4}{3} \times \pi \times r_{min}^3 \tag{32}$$

Then, we compute the parameters of the cell. Since all fluxes in the model are scaled to gram dry cell weight ($gDW$), it is handy to convert the units of cell volume to cell dry biomass. (Canelas et al., 2011) have reported that 1 gDW of cells occupy a volume of roughly 1.7 mL, and this number is relatively stable at different growth rates:

$$V_{gDW} = 1.7\ mL\ gDW^{-1} \tag{33}$$

Even though the volume-dry weight relationship is fixed, it is known that faster-growing *S. cerevisiae* cells are larger. We used experimental data from (Tyson et al., 1979) to compute growth rate $\mu$-dependent cell volume:

$$V_{cell}\ [\mu m^3] = 29.3716 \times \mu + 20.55066 \tag{34}$$

Assuming a spherical cell (close to the natural shape of *S. cerevisiae*), we can compute the radius of such a cell, and, subsequently, its surface area.

$$r_{cell}\ [\mu m] = (\frac{4}{3} \times \frac{1}{\pi} \times V_{cell})^{\frac{1}{3}} \tag{35}$$

$$A_{cell}\ [\mu m^2] = 4\pi \times r_{cell}^2 \tag{36}$$

Following Eqs. 33 and 34, a number of cells in a gram dry weight can be computed:

$$N_{cells} = \frac{V_{gDW} \times 10^{-6}\ [\frac{\mu m^3}{mL}]}{V_{cell} \times 10^{-18}\ [\frac{m^3}{\mu m^3}]} \tag{37}$$

##### 3.2.1 Plasma membrane area constraints

We use the following constraint expression to define the protein capacity in the plasma membrane of the cell. In the model, we assume that 20% of the plasma membrane ($PM$) area is accessible for proteins to occupy ($f_{PM}$). Then, for two specific groups of proteins, transporters of carbon sources ($PMC$) and nitrogen sources ($PMN$), we formulate two separate constraints, assuming these transporters can occupy 9.5% ($f_{PMC}$) and 5.0% ($f_{PMN}$)of plasma membrane surface area, respectively.
First we compute the sum of the molecular weight of the protein complex, formed in the plasma membrane, based on the molecular weight ($MW$) and stoichiometry ($n$) of the proteins, forming the complex:

$$MW_{cplx} = \sum_{i \in cplx} n_i \times MW_i \tag{38}$$

Using Eqs. 30 and 31, the crossection area of this protein complex (per $mmol$ of complex) is computed as follows:

$$A_{mmol\ cplx} = A_{peptide}(MW_{cplx})\ [nm^2] \times 10^{-6}\ [\frac{\mu m^2}{nm^2}] \times 6.022 \times 10^{20}\ [mmol^{-1}] \tag{39}$$

Next, we can compute the surface area the $mmol$ of this protein complex would occupy per single cell (based on the cell number computation from Eq. 37):

$$A_{mmol,sc} = \frac{A_{mmol\ cplx}}{N_{cells}} \tag{40}$$

Finally, we formulate the constraint of available plasma membrane area per single cell:

$$\sum_{i \in compartment} \frac{v_{cplx\ formation,\ i}}{k_{deg} + \mu} \times A_{mmol,sc}\ \leq f_{compartment} \times A_{cell}\ [\mu m^2\ cell^{-1}] \tag{41}$$

Note that here we sum not over protein translation fluxes ($v_{syn}$), but through complex formation fluxes ($v_{cplx\ formation,\ i}$). This is a precaution to make sure that only these proteins, which form complexes, attributed to plasma membrane (the target compartment remains cytosol), are counted. In the mitochondrial volume constraint (see below), sum is also computed not through $v_{syn}$ reactions due to the same issue.

##### 3.2.2 Mitochondrial volume constraint

Similarly to the previously described constraint, we compute the volume of a single peptide and scale it per $mmol$ and per single cell. Since there is no ambiguity of whether the protein should be attributed to mitochondrial capacity or not (see the Eq. 44), we sum over individual proteins rather than their complexes.

$$V_{mmol} = V_{peptide}\ [nm^3] \times 10^{-9}\ [\frac{\mu m^3}{nm^3}] \times 6.022 \times 10^{20}\ [mmol^{-1}] \tag{42}$$

$$V_{mmol,sc} = \frac{V_{mmol}}{N_{cells}} \tag{43}$$

We formulate a mitochondrial volume constraint in the following way, summing the fluxes through protein folding reactions in mitochondria ($v_{folding,\ mito,\ i}$):

$$\sum_{i \in Mito} \frac{v_{folding,\ mito,\ i}}{k_{deg} + \mu} \times V_{mmol,sc}\ \leq V_{mito\ proteins}\ [\mu m^3\ cell^{-1}] \tag{44}$$
